# Membraneless organelles formed by liquid-liquid phase separation increase bacterial fitness

**DOI:** 10.1101/2021.06.24.449778

**Authors:** Xin Jin, Ji-Eun Lee, Charley Schaefer, Xinwei Luo, Adam J. M. Wollman, Tian Tian, Xiaowei Zhang, Xiao Chen, Yingxing Li, Tom C. B. McLeish, Mark C. Leake, Fan Bai

## Abstract

Liquid-liquid phase separation is emerging as a crucial phenomenon in several fundamental cell processes. A range of eukaryotic systems exhibit liquid condensates. However, their function in bacteria, which in general lack membrane-bound compartments, remains less clear. Here, we used high-resolution optical microscopy to observe single bacterial aggresomes, nanostructured intracellular assemblies of proteins, to undercover their role in cell stress. We find that proteins inside aggresomes are mobile and undergo dynamic turnover, consistent with a liquid state. Our observations are in quantitative agreement with phase-separated liquid droplet formation driven by interacting proteins under thermal equilibrium that nucleate following diffusive collisions in the cytoplasm. We have discovered aggresomes in multiple species of bacteria, and show that these emergent, metastable liquid-structured protein assemblies increase bacterial fitness by enabling cells to tolerate environmental stresses.

**One Sentence Summary:** Bacteria use subcellular proteinaceous liquid droplets to survive stress

## Main Text

Liquid-liquid phase separation (LLPS) drives the formation of membraneless compartments in eukaryotic cells, such as P-granules, nucleoli, heterochromatin and stress-granules (*1-5*), enabling concentrations of associated biomolecules to increase biochemical reaction efficiency, and protection of mRNA or proteins to promote cell survival under stress. Compared to eukaryotes, bacterial cytoplasm is more crowded and in general lacks membrane-bound organelles (*6*). Recently, we revealed the existence of bacterial aggresomes, subcellular collections of endogenous proteins that appear as dark foci in the cell body (*7*), which are dynamic, reversible structures that form in stressed conditions and disassemble when cells experience fresh growth media (*7*) (Fig. S1A). Here, we exploit tandem experiment and multiscale modeling to determine the molecular biophysics of their spatial and temporal control, and deduce their crucial role in bacterial fitness.

We initially screened three proteins (HslU, Kbl and AcnB) as biomarkers, according to our previous analysis of their abundance in aggresomes, and labeled each with fluorescent protein reporters at chromosome (Tables S1-3). All three accumulated in aggresomes, though with different degrees of colocalization to dark foci in brightfield images (Fig. S1B). We overexpressed HokB protein, a toxin causing cellular ATP depletion, to trigger aggresome formation in *E. coli* (*8*) (Fig. S1C). Monitoring the proportion of cells containing distinct fluorescent foci from time-lapse epifluorescence microscopy indicated different clustering dynamics with respect to ATP depletion (Fig. 1A). HslU, which had the highest colocalization to dark foci and was the most rapid responder with respect to ATP depletion, showed distinct fluorescent foci in 50% of cells after 60 mins HokB induction, followed by Kbl (requiring 150 mins), while the formation of AcnB foci was significantly slower. Dual-color fluorescence imaging confirmed the colocalization between HslU, Kbl and AcnB and their differing dynamics of incorporation into aggresomes (Fig. 1B, Fig. S1D).

**Fig. 1.**
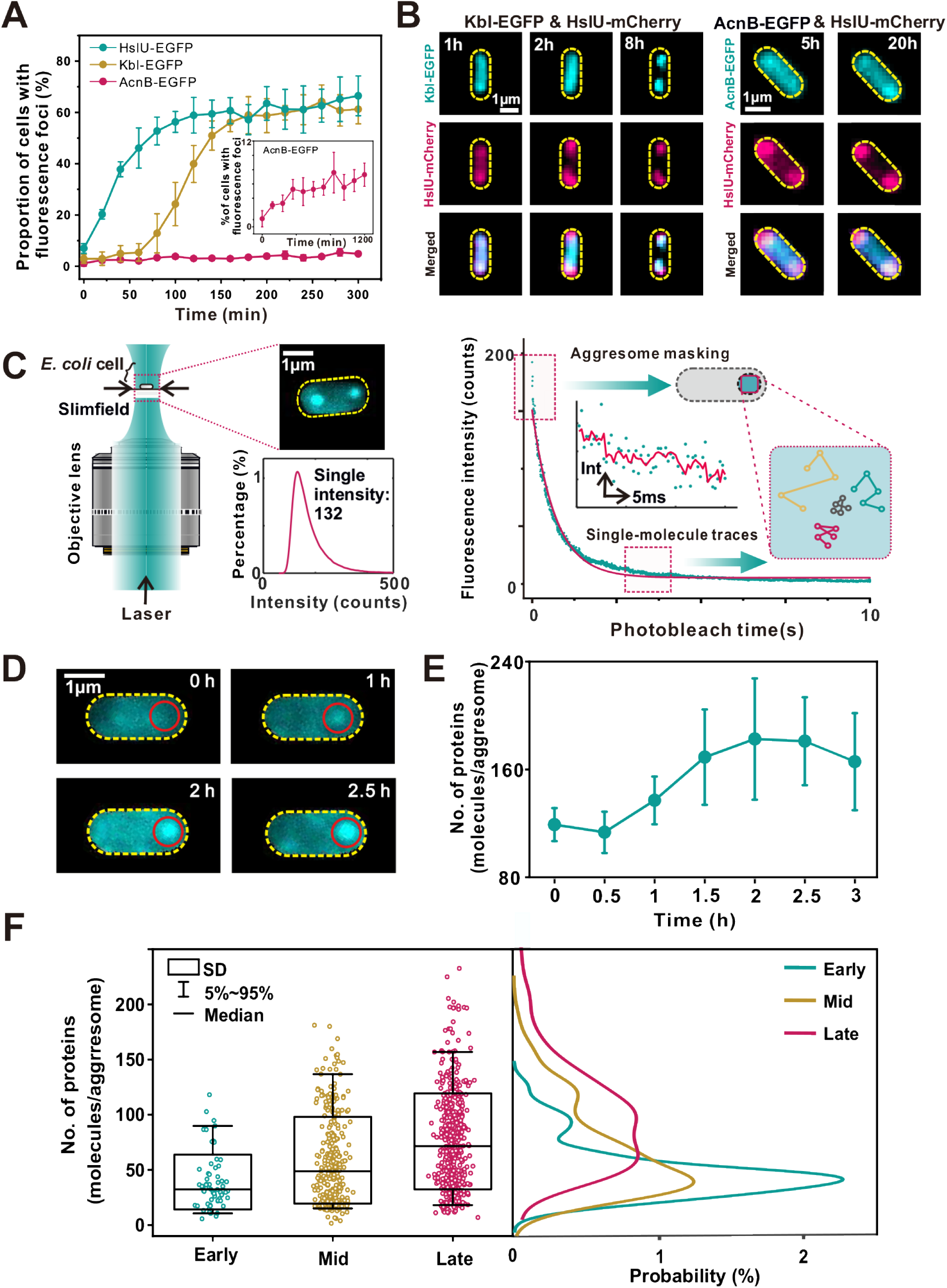
Aggresome formation in bacteria. (A) Proportion of cells showing fluorescent foci as a function of HokB induction time. (B) Fluorescence images of dual-labeled strains indicating protein-dependent dynamics of cluster formation. (C) Left: Schematic of Slimfield microscopy used for rapid super-resolved single-molecule tracking. The value of single intensity (132 counts on our camera detector) is the characteristic integrated intensity of a single fluorescent EGFP molecule. Right: For detecting aggresomes, we used a five average image at the start of the photobleach of each image acquisition. For tracking single molecules, we looked into the end of photobleach processes. (D) Fluorescence images for strain HslU-EGFP during aggresome formation (red circle). (E) Number of HslU-EGFP per aggresome from (D), SD error bars. (F) Left: box plots for number of HslU-EGFP per aggresome at different time points. Number of aggresomes N=62, 306, and 490 for early, mid, and late respectively. Right: Kernel density estimation (*9*) of the number of EGFP molecules per aggresome at the different HokB induction stages.

Since HslU showed the highest abundance and rate of response following ATP depletion we used it as the best biomarker from the three candidates to explore spatiotemporal features of aggresome formation with rapid super-resolved single-molecule tracking (Fig. 1C). We applied step-wise photobleaching analysis (*10*) to these data to determine the number of HslU-EGFP molecules visible inside and outside aggresomes in individual live cells (Fig. 1D). These data showed that the mean HslU-EGFP stoichiometry within aggresomes with respect to HokB induction time increases initially and then saturates, indicating demixing of HslU-EGFP from the surrounding cytoplasm in response to ATP depletion over a timescale of 1-3 hrs (Fig. 1E), that we categorized into ‘early’ (0.5 hr after induction, N=31), ‘mid’ (1 hr after induction, N=130), and ‘late’ (2-3 hrs after induction, N=209) stages (Fig. 1F). 3D structured illumination microscopy (3D-SIM) was used to determine the morphology of aggresomes at different stages (Fig. S1E). The mean aggresome diameter is 178 ± 6 nm (± SE, early stage, number of aggresomes=100), to 242 ± 8 nm (mid stage), and then 309 ± 5 nm (late stage), with associated sphericity of 0.817 ± 0.005 (early stage), 0.841 ± 0.005 (mid stage), and then 0.927 ± 0.004 (late stage).

To investigate the patterns of mobility for HslU in bacterial cytoplasm, we photobleached the majority of HslU-EGFP and tracked the movement of fluorescent foci with only a few HslU molecules (Fig. S2A). We determined the apparent diffusion coefficient (*D*) of foci from the initial gradient of the mean square displacement (MSD) with respect to tracking time (Fig.2A, showing the spatial mapping of representative diffusive trajectories of HslU-EGFP proteins in the bacterial cytoplasm). To determine the molecular mobility of HslU inside aggresomes we tracked the fluorescent foci with only a single molecule, as confirmed by brightness and circularity (Fig. S2B and S2C). The mean diffusion coefficient for single molecule HslU-EGFP (*D*_*g*_) across a population of cells decreased from 0.32 ± 0.05μm^2^/s (± SE, early stage, number of tracks N=183), to 0.24 ± 0.04 μm^2^/s (mid stage, N=176), and then 0.19 ± 0.02 μm^2^/s (late stage, N=303) (Fig. 2B). Tracking single HslU-EGFP also revealed MSD maxima showing an aggresome diameter of up to a mean of 305 nm at late stage (Fig. S2D), which is consistent with 3D-SIM measurements.

**Fig. 2.**
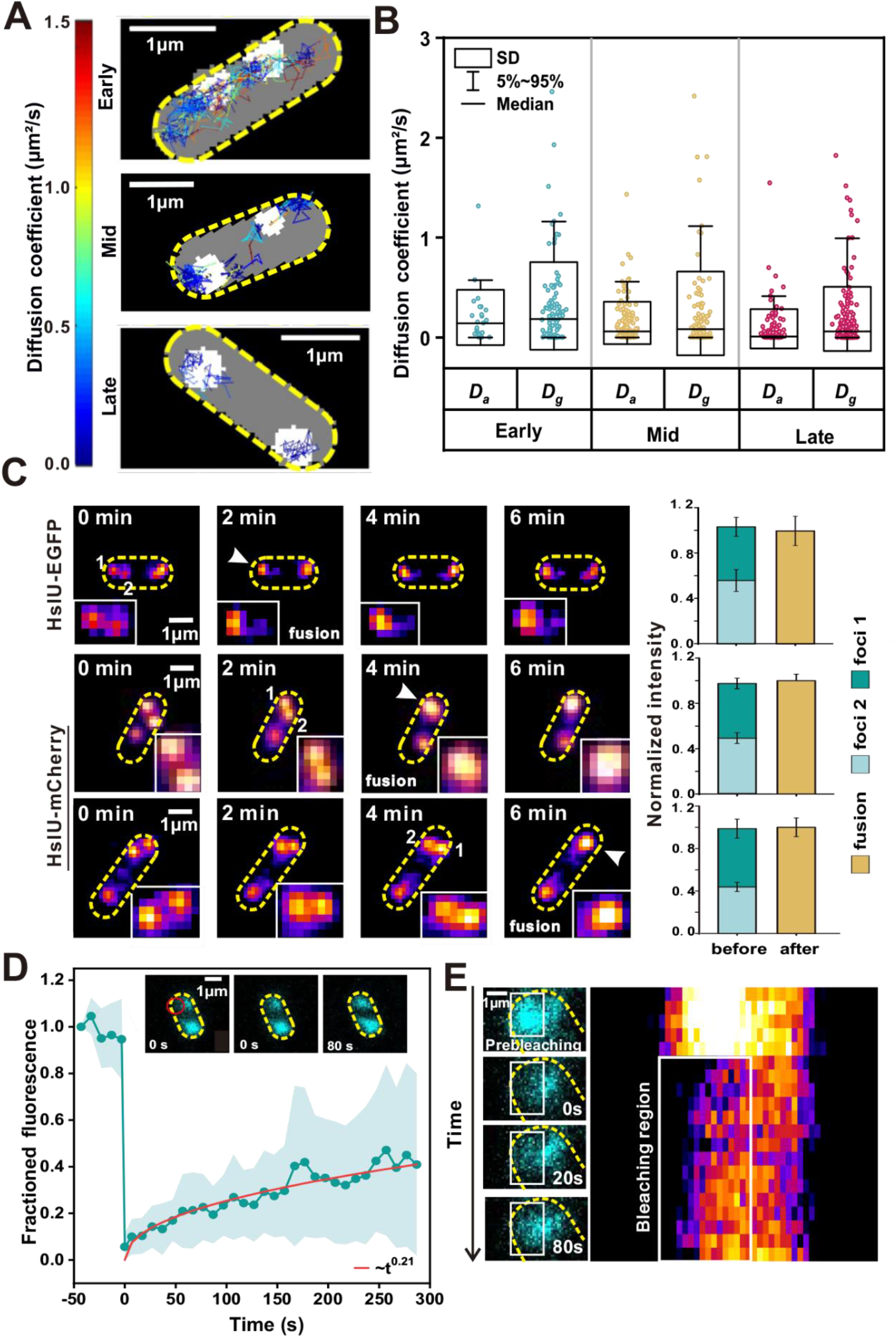
Aggresome formation occurs through LLPS. (A) Representative tracking of HslU-EGFP foci during aggresome formation, color corresponding to diffusion coefficient (left bar). (B) Box plots of *D*_*g*_ and *D*_*a*_ of HslU-EGFP at different time points. (C) Left: Representative epifluorescence images of two aggresomes fusing. Right: Total fluorescence intensity of aggresomes before and after fusion, error bar represents the SD pixel noise. (D) Mean fluorescence recovery curve (cyan) defined from half region of an aggresome indicating a mean recovery half time of 50 ± 9 secs and a heuristic power-law fit (red: ∼t^0.21^), with inset half-FRAP images of an aggresome showing fluorescence recovery, position of focused FRAP laser waist indicated (red). Error bounds show SD (number of aggresomes N=29, each from a different cell). (E) Zoom-in of aggresome showing fluorescence recovery with kymograph (right heatmap).

We then corrected the apparent diffusion coefficient of these single-molecule tracks by subtracting the bulk diffusion measured for the whole aggresome itself (*D*_*a*_), which was obtained by tracing the movement of the centroid of the aggresome, to yield the diffusion estimates for individual HslU molecules relative to its confining aggresome. The mean values of *D*_*a*_ were 0.20 ± 0.06 μm^2^/s (early stage), 0.15 ± 0.02 μm^2^/s (mid stage), and 0.09 ± 0.01 μm^2^/s (late stage) (Fig. 2B). To check that these *D*_*a*_ values are biologically reasonable, we modeled the aggresome diffusion (*D*_*a*_) using the Stokes-Einstein relation for viscous drag associated with a sphere of diameter 305 nm that indicated a local cytoplasmic viscosity of 16.6 cP (1 cP = 1 mPa·s) (Table S4), a value broadly consistent with previous estimates on live *E. coli* (*11, 12*). The mean values of *D*_*g*_*-D*_*a*_were 0.12 ± 0.07 μm^2^/s (early stage), 0.09 ± 0.05 μm^2^/s (mid stage), and 0.10 ± 0.03 μm^2^/s (late stage), contrasted against 0.05 ± 0.02 μm^2^/s (greater than zero due to finite photon sampling) in separate experiments using identical imaging on single EGFP *in vitro* immobilized via antibodies to the glass coverslip surface (*13*) (Fig. S2E). Importantly, therefore, single HslU-EGFP inside aggresomes diffuse faster than immobilized EGFP at all stages. HslU molecules are thus definitively mobile inside aggresomes, consistent with a liquid state.

A common feature of LLPS is fusion of droplets to form larger, spherical compartments. To investigate this characteristic in aggresomes, we performed time-lapse epifluorescence microscopy at 2 min intervals to capture fusion events, using EGFP and mCherry reporters of HslU to ensure that this was not uniquely associated with just one type of fluorescent protein. As shown (Fig.2C and Fig. S3A) foci fusion often occurred at early- or mid-stage aggresome formation, with the fluorescence intensity of merged compartments tallying with the sum of intensities of individual foci to within measurement error. Also, the circularity of the merged aggresome peaked at approximately 1 (Fig. S3B). These observations suggest that larger aggresomes may result from fusions of smaller HslU droplets, and support the hypothesis that LLPS drives bacterial aggresome formation.

To further test the liquid nature of aggresomes, we implemented FRAP measurements. By focusing a laser laterally offset approximately 0.5 µm from an aggresome center it was possible to photobleach EGFP content within approximately one spatial half of the aggresome, while leaving the other half intact, denoting this “half-FRAP”. We then measured the aggresome fluorescence intensity at 10 sec intervals for up to several hundred seconds afterwards (Movie S1). Fluorescence in the bleached region recovered with a half time 50 ± 9 sec (±SD) to approximately 35% of the initial intensity at steady-state (Fig. 2D, kymograph shown in Fig. 2E). Similarly, we observed fluorescence loss in photobleaching (FLIP) for the initially unbleached half with increasing time after the focused laser bleach (Fig. S3C). Taken together, these results reveal that HslU-EGFP undergoes dynamic turnover within the aggresome over a timescale of tens of seconds.

To investigate turnover of HslU-EGFP between aggresomes and the surrounding cytoplasm we performed “whole-FRAP” for which a focused laser was centered directly on an aggresome, enabling us to photobleach all its EGFP contents while leaving the bulk of the remaining fluorescence in the cell, including that for other aggresomes, intact. We saw that the fluorescence of bleached aggresomes recovered with a half-time of 32 ± 18 sec, almost twice as rapidly as half-FRAP, though the recovery fraction of whole-FRAP (∼10%) is much lower than that of half-FRAP (Fig. S3D). We also noticed that the rate of loss of fluorescence for the equivalent whole-FLIP was greater if we measured the loss from the whole cell area outside a bleached aggresome as opposed to just the fluorescence from a second aggresome localized to the opposite pole of the same cell, suggesting that short timescale turnover is limited to the local vicinity outside an aggresome. The whole-FRAP measurement indicates that there is a turnover of HslU molecules inside and outside aggresomes.

We developed an Individual-Protein-Based Model (IPBM) (*14, 15*) to interpret the experimental observations by simulating collective dynamics of aggresome formation, protein turnover and dynamics as LLPS under thermal equilibrium (Fig. 3A). From our previous work, we know aggresome formation involves many types of proteins (*7*). Here we model this ensemble effect using an effective field approximation, denoting non-biomarker proteins that participate in aggresome formation as a general protein A while a specific aggresome biomarker protein is denoted B. In the first instance we used the experimental results from HslU-EGFP as protein B to establish a value for the interaction energy between A and B. We are also able to interpret the results from aggresome proteins Kbl and AcnB using the same model but with different interaction energies (Fig S5H).

**Fig. 3.**
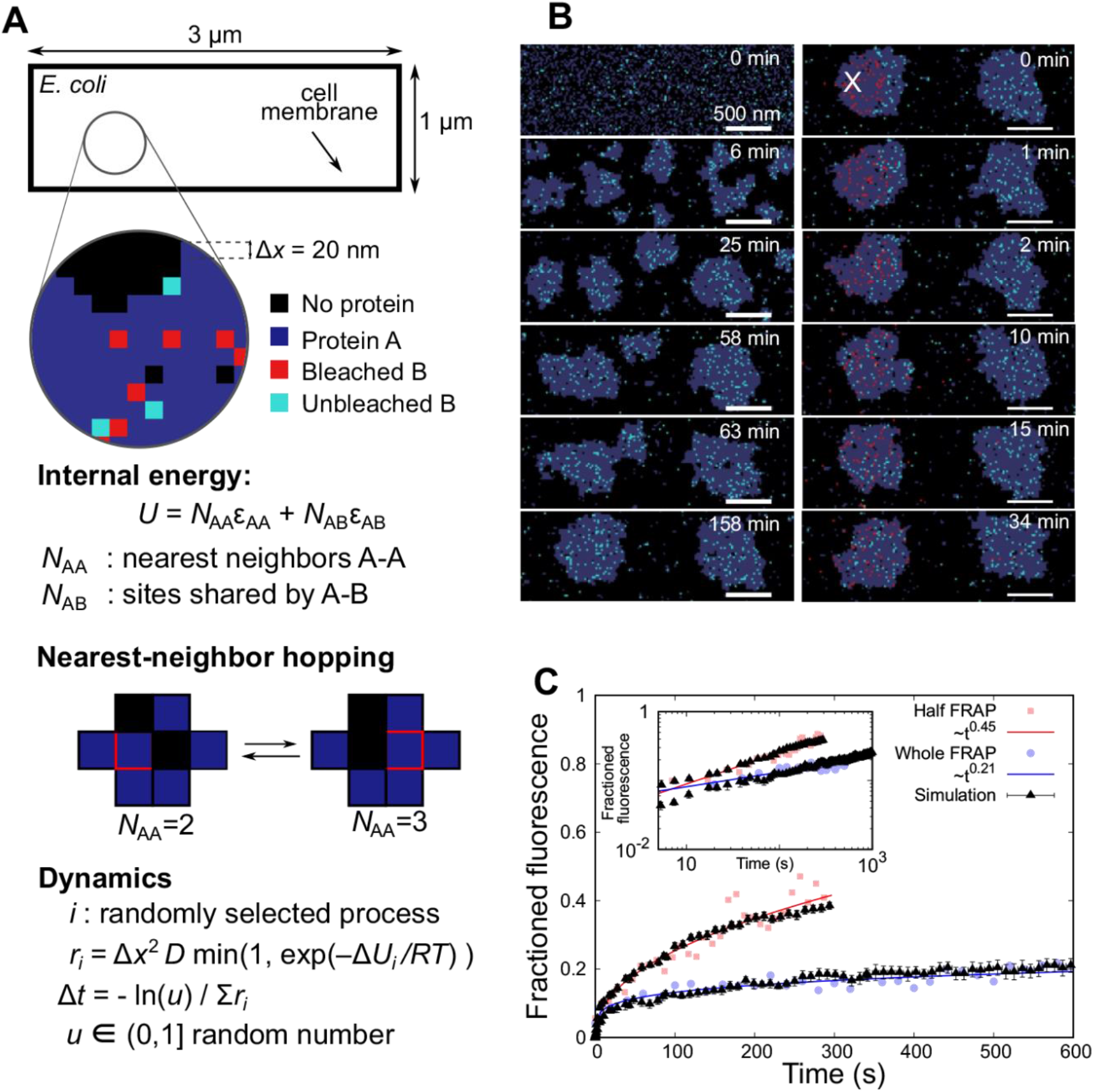
Simulating aggresomes using an Individual-Protein-Based Model. (A) The cell is modeled as a 3×1 µm 2D rectangular lattice with regular 20 nm lattice spacing. Each lattice site can be either vacant, occupied by a single protein A (gray) or B (cyan), or by two proteins that are interacting (A and B). Protein A represents a general component in the aggresome that drives LLPS through an attractive nearest-neighbor interaction energy (optimized here to ε_AA_=2.2 *k*_*B*_*T*) with protein B a fluorescently-labeled aggresome biomarker (such as HslU). The total internal energy is *U = N*_AA_*ε*_*AA*_+ *N*_AB_ *ε*_*AB*_, with *N*_AA_the number of A-A nearest-neighbor pairs, and *N*_AB_ the number of sites shared by proteins A and B. Dynamics are captured using a stochastic Monte Carlo algorithm in which proteins move to nearest-neighbor sites with rate *r* dependent on diffusivity *D*. (B) Simulations predict early-stage LLPS and coarsening by both Ostwald ripening and droplet fusion, resulting in two metastable droplets. Droplets form near cell poles and eventually merge but are stable within the experimental time scales. Proteins in a circular region of radius 350 nm are photobleached (red) after a time of approximately 140 minutes to simulate FRAP. (C) By quantifying the number of protein B molecules entering the photobleaching region we obtain simulated FRAP curves that obey scaling relationships that are in good quantitative agreement with whole- and half-FRAP experiments, here shown for HslU-EGFP (*R*^*2*^ goodness-of-fit values of 0.756 and 0.917 respectively). 2,350 protein A molecules used equivalent to a volume fraction of 31.3%, with 400 protein B molecules.

The interactions and dynamics were modeled on a 2D rectangular lattice of 1×3 µm^2^ (representing a geometrically simplified single *E. coli* cell) with grid spacing Δ*x*=20 nm. Each lattice site may be vacant or occupied by a single protein A or B, or both. The internal energy is:

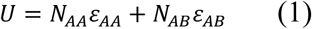

where *N*_AA_ is the number of nearest-neighbor pairs of A, *ε*_AA_ the corresponding pair-interaction energy, while *N*_AB_ is the number of sites shared by the A and B proteins and *ε*_AB_ is the interaction energy between A and B. We found that *ε*_AA_=2.2 *k*_B_*T*, which is above the critical value of ∼1.8 *k*_B_*T* for LLPS (*16*), resulted in the best agreement to the experimental data. Within the framework of this model, it is expected that LLPS triggered by ATP depletion may be modeled by a change of the interaction parameter from *ε*_AA_<1.8*k*_B_*T* to *ε*_AA_>1.8*k*_B_*T*.

We modeled the molecular dynamics using a (stochastic) Monte Carlo process where proteins hop to nearest-neighbor sites with an attempt frequency (*17*)

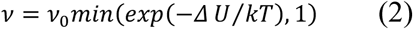

where
ν_0_=*D*/Δ*x*^2^ depends on the bare diffusivity *D* in the absence of interactions, and Δ*U* is the energy difference due to the hop of a protein into a new local environment. We chose to model, at the interface a hop of B into or out of the aggresome is associated with a net diffusivity (*D*_B,in_+*D*_B,out_)/2. The time step for each hop depends on the sum, *S*, of all possible rates in Eq. (2) as Δt = –ln(*<u>u</u>*)/ *S*, with 0<*<u>u</u>≤*1 a random number.

We generated representative LLPS behaviors (Fig. 3B and Movie S2), showing early-stage small clusters that coarsen according to a 1/3 power scaling with time as expected from Ostwald ripening and Brownian coalescence (Fig S4A-D) (*18*). The key observation addressed by the time-dependent behavior is the qualitative feature of two droplets of comparable stoichiometry to experimental data forming at cell poles (Fig. 3B and Movie S2) over a timescale of 1-2 hours relevant to HokB induction times that only merge at much longer time scales. These model- interpreted observations suggest: (i) the long time scale does not necessarily point at the need to overcome a high free-energy to nucleate aggresomes (*19*), and (ii) that the observation of two aggresomes is a consequence of the elongated topology of the cell without a need of any activated processes; although the nucleoid in the center of the cell (absent in our simulations) may further stabilize the two-droplet morphology (Movie S3).

The same model without change of parameters was able to reproduce simultaneously both the whole- and half-FRAP experiments (Fig. 3B-C). Here, we simulated LLPS using IPBM for approximately 140 mins prior to photobleaching a sub-population of protein B molecules localized to a laser bleach zone (approximated as a circle of radius 350 nm), after which the IPBM simulation continued and we monitored recovery as the number of unbleached protein B molecules in the laser bleach zone. In half-FRAP, fast recovery (∼1 min) is governed by diffusion of unbleached B inside aggresomes, absent in whole-FRAP. At ∼2 mins, recovery is more dependent on diffusion of unbleached B from the cytoplasm immediately surrounding aggresomes. At longer times, B from the opposite pole can reach the bleach zone and influence fluorescence recovery. We found that the experimental FRAP data for other aggresome proteins of Kbl and AcnB tried in our initial screen showed half-FRAP recovery to be consistent with a 1/2 power law (Fig. S5E). Therefore, we collapse all experimental and simulation data onto half-FRAP master curves (Fig. S5F-H). The differences in the offset of the whole-FRAP curves for the three types of proteins (Fig. S5H) seems to originate from differences in the binding energies of the respective proteins to the aggresome (Fig. S5F) and/or variations in their mobility in the cytoplasm (Fig. S5G). Our straightforward theoretical framework demonstrates that relatively simple protein physics – energetic interaction of diffusing proteins under local thermal equilibrium characterized by a simple and coarse-grained pairwise interaction energy – accounts not only for the conditions of aggresome formation in the first place, but also for complex, emergent biological features in their spatial localization and their dynamics and kinetics of molecular turnover.

Finally, we investigated if aggresomes confer physiological advantages to bacteria. We noticed that aggresomes are not unique to *E. coli*; by prolonged stationary culturing, we observed them in all eight of the other Gram-negative species we investigated (Fig. S6A). We performed large-scale screening of small chemical compounds and the *E. coli* gene knock-out library, and found three representative conditions, comprising media supplementation with MOPS (*20*) and deletions *ΔsdhC* and *ΔnuoA (21*), that disabled aggresome formation after prolonged stationary culturing (Fig. 4A-B).

**Fig. 4.**
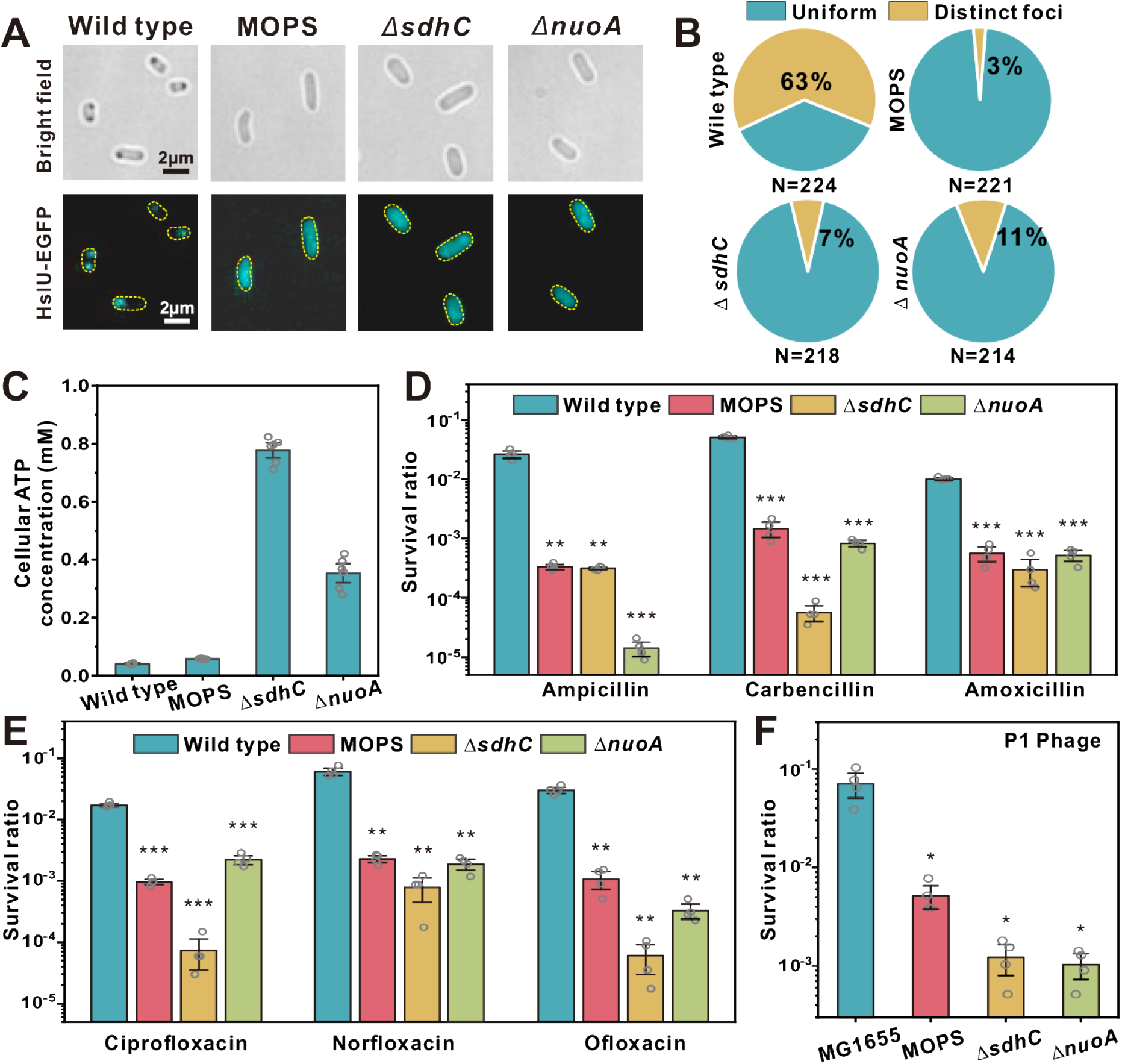
Aggresome formation promotes cellular survival under fierce stresses. (A) Brightfield and fluorescence images of MG1655, MG1655 (MOPS), *ΔsdhC* and *ΔnuoA*, (B) proportion of cells showing HslU-EGFP foci, and (C) average cellular ATP concentration in different strains, after 24 hrs culture. Cell survival rate (log scale) after 4 hrs (D) β-lactam (E) fluoroquinolone, and (F) P1 phage infection, Multiplicity of Infection (MOI)=100. (Unpaired Student’s t-test against wild type, error bar indicates SE, ^*^p value < 0.05; ^**^p value < 0.005; ^***^p value < 0.0005.)

MOPS has been reported as an osmolyte and protein stabilizer (*20*); in a wild type strain it penetrates the cell and stabilizes proteins from aggregation even if cytoplasmic ATP is depleted (*20, 22*). Both *sdhC* and *nuoA* encode genes for the respiratory chain, where *sdhC* encodes the membrane-anchoring subunit of succinate dehydrogenase, and *nuoA* encodes NADH-quinone oxidoreductase subunit A. In respiration-impaired mutants, *ΔsdhC* and *ΔnuoA* strains, aggresome formation was disabled by inhibiting intracellular ATP decrease (Fig. 4C).

We compared the survival difference between wild type, knockout strains *ΔsdhC* and *ΔnuoA*, and wild type supplemented with MOPS after stationary culturing. Interestingly, we observed that under various antibiotic treatments, the wild type strain with successful aggresome formation showed a higher persister ratio in comparison to strains in which aggresome formation was inhibited (Fig. 4D-E), indicating an increased fitness. We observed the same survival advantage in wild type when challenging bacteria with phage invasion (Fig. 4F). We also defined the survival ratio of these strains in early stationary phase, when no strains exhibited aggresomes, and found that all the strains have similar tolerance in early stationary phase (Fig. S6C-E). Taken together, we conclude that aggresome formation through LLPS promotes bacterial survival under a range of fierce stresses. This is consistent with our previous hypothesis (*7*): the formation of aggresomes sequester numerous proteins which are vital for cellular function, leading to shutdown of different biological processes and the cell enters a dormant state.

The formation of certain small biological condensates in bacteria has recently been ascribed to LLPS (*23, 24-25*). Here, we have established that aggresomes are membraneless liquid droplets hundreds of nm in diameter that phase separate following clustering of diffusing proteins under thermal equilibrium in the bacterial cytoplasm. Our modeling indicates no requirement of external free energy for aggresome formation, for example from ATP hydrolysis as proposed recently for *parS*/ParB liquid condensates in *E. coli* (2*5*), but a dependence on ATP cellular concentration. A possible explanation could lie in recent work which demonstrated that ATP not only energizes processes inside cells but also acts as a hydrotrope to increase specific protein solubility (*26*); decreasing cellular ATP may thus act to favor LLPS. This method of regulation using ATP might be an important and recurring theme in several other biological processes that utilize LLPS.

## Acknowledgments

**Funding:** This work was financially supported by the National Natural Science Foundation of China (No. 31722003, No.31770925) to FB, the Engineering and Physical Science Research Council (EPSRC) (EP/T002166/1, EP/N031431/1) to M.C.L., Biotechnology and Biological Sciences Research Council (BB/P000746/1, BB/R001235/1) to M.C.L. and J.-E.L., Royal Society (IEC\NSFC\191406) to M.C.L., the EPSRC (EP/N031431/1) to C. S. and T. C. B. M.

**Author contributions:** F.B., M.C.L., X.J., J.-E.L. conceived the study; X.J., J.-E.L., X.L., T.T., X.Z. performed the experiments; J.-E.L., X.J., C.S. led the data analysis with help from A.J.M.W.; C.S. led the modeling and theoretical study; X.J., J.-E.L., C.S., F.B., M.C.L. wrote the manuscript with inputs from all authors. We thank the York Bioscience Technology Facility for assistance with confocal microscopy. We thank National Center for Protein Sciences at Peking University in Beijing, China, for assistance with 3D-SIM imaging.

**Competing interests:** The authors declare no competing interests.

**Data and materials availability:** All data is available in the main text or the supplementary materials

