## supplementary methods for "Membraneless organelles formed by liquid-liquid phase separation increase bacterial fitness"

**This PDF file includes:**

Materials and Methods
Supplementary Text
Figs. S1 to S6
Tables S1 to S4
Captions for Movies S1 to S3
References

**Other Supplementary Materials for this manuscript include the following:**

Movies S1 to S3

### Materials and Methods

<u>Bacterial Strains, Phage and Plasmid Construction</u>

All strains used in this study are indicated in Table S1, plasmids in Table S2, and primers in Table S3. Wild-type strain MG1655 was a gift from Yale Genetic Stock Center. P1 phage was a gift from Dr. Xilin Zhao, Xiamen University, China. Strains containing chromosomal *geneX-egfp/mcherry* translational fusion or single geneX knockout mutants were constructed by λ red mediated gene replacement (*27*). For fluorescent protein fusion strains: the targeted fluorescent protein (EGFP or mCherry) fragment was amplified and inserted to replace the stop cassette of the selection-counter-selection template plasmid (pSCS3V31C). Then the linker-FP-Toxin-CmR fragment was amplified from the template plasmid with homology arm complementary to the flanking sequences of the insertion site on the chromosome, before being transformed into electrocompetent cells with induced recombineering helper plasmid (pSIM6) (*28*). After 3-5 hrs recovery, transformed cells were plated on selection plates containing chloramphenicol (25 μg/mL). The Toxin-CmR cassette was then removed from the chromosome by another round of λ red mediated recombination using a counter selection template. Finally, cells were plated on counter selection plates containing rhamnose to activate the toxin (*29*). For knockout strains, we used the Keio cassette flanked by FRT sites in the Keio collection (*30*) to replace the certain gene on the chromosome.

The pBAD::*hokB* plasmid was transformed by electroporation into certain strains to generate *hokB* overexpressing strains. The *hokB* PCR product was amplified from MG1655 and inserted into pBAD/myc-His A vector at *Nco I* and *Hind III* sites by the *in vitro* Gibson Assembly method. The ampicillin resistant gene was replaced with chloramphenicol resistance cassette. All strains were grown in Luria Broth (LB). We determined the cell doubling times of all strains generated to compare against their wild type equivalents (Table S1).

### Microscopy

#### *Time-lapse microscopy*

Standard brightfield and epifluorescence imaging were performed on an inverted microscope (Zeiss Observer Z1, Germany). Illumination was provided by solid-state lasers (Coherent OBIS, USA), at excitation wavelengths 488 nm for EGFP and 561 nm for mCherry. The fluorescence emission signals of cells were imaged onto an EMCCD camera (Photometrics Evolve 512, USA). GFP ($\lambda_{em}$=500-550 nm) and mCherry ($\lambda_{em}$=604-640 nm) Zeiss filter sets were selected for each fluorophore according to their excitation and emission spectra.

Aggresome formation time-lapse imaging was performed using the Flow Cell System (Bioptechs FCS2, USA). Cells from the early stationary phase were washed with PBS using a centrifugation and were tipped on a gel-pad containing 2% low melting temperature agarose and 0.1% arabinose. Cells were then observed under brightfield and epifluorescence illumination at 37 ºC.

Home-written MATLAB software was used to detect fluorescent foci in time-lapse images. Cells were segmented according to brightfield images. Then, the fluorescence distribution inside the cells was analysed to determine whether there is a fluorescent foci. We used a 3×3 pixel (0.47×0.47 $\mu m^2$) kernel search for distinct fluorescent foci pixel-by-pixel inside each cell. If the foci met the two following conditions at the same time, we defined it as a fluorescent 'focus'. First, the mean fluorescence of the focus is higher than 1.5 times the mean fluorescence intensity of the whole cell. Second, the mean fluorescence of the foci is higher than 1.5 times of the mean local background intensity located in a 2 pixel ring area around a candidate focus. For brightfield black foci detection, cells were segmented first (excluding the white edges around each cell body image).Then we detected the black foci in the cells based on the gray value and area. If the gray value of a pixel is less than 95% of the average gray value of the whole cell, the pixel will be listed as a candidate of black foci. After the connected domain measurement, if the area is larger than 4 pixels, it will be defined as a black focus. The colocalization analysis was performed by analyzing the Pearson correlation coefficient of EGFP and mCherry fluorescent foci using Fiji plugin JACoP (*31*).

*Slimfield microscopy*

Microscopy was performed utilizing narrow epifluorescence excitation of 10 μm full width at half maximum (FWHM) in the sample plane to generate a Slimfield excitation field in the sample plane (*32*). EGFP was excited by a linearly polarized 488 nm wavelength 50 mW laser (Coherent OBIS, USA) attenuated to approximately 7 mW at the sample with average excitation intensity 4.6 kW/cm$^2$. To generate a circularly polarized beam, the laser was directed through a λ/4 waveplate and then to a dual-pass green/red dichroic mirror centered at long-pass wavelength 560 nm. Fluorescence emissions were captured by a 1.49 numerical aperture (NA) oil immersion objective lens (Nikon Apo TIRF, Japan) and passed through a bandpass emission filter with a 25 nm bandwidth centered at 525 nm (Chroma Technology, USA). Fluorescence emissions were imaged onto a Photometrics 95B CMOS camera, magnified to 50 nm/pixel. For the sample imaging protocol, a brightfield image was first acquired for 2 frames, and then we acquired EGFP images by the 488 nm excitation laser beam using a 5 ms exposure time per frame for 2,000 consecutive frames. For aggresome formation, the sample preparation used was the same as that in time-lapse microscopy.

*3D Structure Illumination Microscopy (3D-SIM)*

3D-SIM images were acquired on a DeltaVision OMX SR imaging system (GE Healthcare, USA) equipped with a 100× oil-immersion objective (NA 1.49) and EMCCD, which achieves imaging of samples at approximately 120 nm lateral and 300 nm axial resolution. Laser lines at 488 and 561 nm were used for excitation. The microscope was routinely calibrated with 200 nm fluorescent microspheres. Serial z-stack sectioning was carried out at 125 nm intervals. Images were reconstructed with the softWoRx 5.0 software package. Reconstructed SIM data was further analyzed using software Imaris 9.6.0. The diameter of an aggresome is the equivalent spherical diameter (the diameter of a sphere of equivalent volume as the aggresome). The sphericity is the ratio of the surface area of a sphere (with the same volume as the aggresome) to the surface area of the aggresome. For reference, the sphericity of fluorescent microspheres is 0.973±0.013.

Cells from the early stationary phase were washed and resuspended with PBS. Arabinose (0.1%) was added to induce aggresome formation. Cells were collected at different time points (0.5 h for early, 1 h for mid and 3 h for late) and fixed in 2% PFA solution for 15 mins at room temperature and 30 mins at 4°C.

*Fluorescence Recovery after Photobleaching (FRAP) experiment*

FRAP was performed using a Zeiss LSM 710 confocal microscope with a 63× oil immersion objective (NA 1.4) (Zeiss Plan-Apochromat DIC M27, Germany). For half-FRAP experiments, a circular region of interest (ROI) of about 0.65 μm diameter in a half region of one aggresome (roughly 0.5 μm away from the aggresome's edge) was bleached using a laser focus with 488 nm wavelength generated by an Argon ion laser using a scanning time (i.e. bleaching time) equal to

1.62 frames per second (equivalent to 620 ms). The half region of one aggresome was bleached followed by the monitoring of the time course (every 10 seconds, 35 cycles, the first 5 cycles was the pre-bleaching sequence, and the remaining 30 cycles was the post-bleaching sequence) of fluorescence recovery. For whole-FRAP experiments, a circular ROI of about 1 μm diameter in one aggresome was bleached using the same laser with a scanning time (i.e. bleaching time) equal to 1.62 frames per second (equivalent to 620 ms). One selected aggresome was bleached followed by the monitoring of the time course (every 20 seconds 35 cycles), the first 5 cycles was the pre-bleaching sequence, and the remaining 30 cycles was the post-bleaching sequence) of fluorescence recovery. The emission detection wavelength ranged from 490 nm to 585 nm. The fluorescence recovery in the bleached region of interest was recorded by the commercial software (Zen Black) that controlled the microscope.

Foci Detection, Tracking, Stoichiometry and Aggresome Masks

Fluorescent foci were automatically detected using a custom program written in MATLAB (*33*) enabling estimation of the number of proteins per aggresome and the apparent diffusion coefficients for the whole aggresome and individual aggresome proteins. The detection and tracking software objectively identifies candidate fluorescent foci above a signal-to-noise ratio (defined as SN) fixed at 0.4 by a combination of pixel intensity thresholding and image transformation to yield initial approximations for the intensity centroid to the nearest pixel. The sub-pixel refinement to the centroid of each focus was then determined using iterative Gaussian masking. The intensity was defined as the summed pixel intensity inside a 5 pixel radius circular ROI corrected for the local background taken as the mean pixel intensity included in a 17×17 square ROI centered on the centroid but excluding the inner circular ROI. Each candidate focus was then fitted subsequently to an unconstrained 2D Gaussian fit to determine the separate σ widths in x and y relative to the camera detector. The circularity of that focus was then defined as $\sigma_x/\sigma_y$.

Characteristic intensity distributions of single EGFP molecules (Figure 1C) were rendered as Kernel density estimation (KDE) (*9*). Distributions were determined from the tracked foci intensity distributions from the end of the photobleach process confirmed by overtracking foci beyond their bleaching to generate individual steps of the characteristic intensity. For this, we collated all the foci after 1/3 bleaching. By doing that, only single fluorophore molecules are detected, that enabled estimation of the characteristic brightness of a single EGFP molecule in a living cell. This was qualitatively compared to *in vitro* estimates from surface immobilized EGFP (Fig S2F), which agrees within 89 %.

Stoichiometry was determined by implementing a linear-fit to the first four intensity values of foci in each track, using the straight line going to just one frame ahead from the first frame of each track to define the initial intensity of each track and dividing this by the characteristic intensity of single EGFP.

Aggresome masks were defined from Slimfield image data by using the same spot detection algorithm as above but instead using a 5 frame average at the start of the photobleach of each image acquisition, resulting in comparable numbers of aggresomes detected per cell as those from detection of the distinct fluorescent foci from slower sampled epifluorescence microscopy. These aggresome foci were then fitted with a 2D radially symmetrical Gaussian function. A circular mask for each aggresome was then set with the center at the Gaussian centroid using a diameter of 1.5× the sigma width of the fitted Gaussian, resulting in a range of diameters across all aggresomes detected of 3-12 pixels.

Mobility Analysis

The two-dimensional apparent diffusion coefficient relative to the camera detector for each track was calculated from the gradient $G$ from a linear fit to first four data points 4 mean square displacement (MSD) values with respect to tracking time interval, i.e. equivalent to time interval values of 5, 10, 15, and 20 ms, with the fit intercept with the zero-time axis constrained to pass at the through $L^2$ where $L$ is the equivalent two dimensional localization precision estimated previously to be approximately 40 nm (*32*). $D$ was then determined as $G/4\Delta t$, where $\Delta t$ =20 ms.

*Dg* denotes the values of apparent diffusion coefficient associated with tracks that were associated with just a single EGFP molecule. We confirmed that these were single-molecule tracks by looking at their stoichiometry which is 1 to within experimental error (Figure S2B). The *Dg* was calculated by fitting the first 4 MSD values from the foci tracks, as above, found at the end of photobleach process. The mean values of *Dg* were 0.32 ± 0.05 µm²/s, 0.24 ± 0.04 µm²/s and 0.19 ± 0.03 µm²/s for early-, mid-, and late-stage respectively.

We then corrected the apparent diffusion coefficient of these single EGFP molecule tracks inside aggresomes by subtracting the diffusion due to the associated aggresome (*Da*). *Da* was calculated by fitting the first 4 MSD values from the foci tracks found within just the first five image frames of the start of laser illumination (i.e. the image frames used to generate the aggresome mask). The mean values of *Da* were 0.20 ± 0.06 µm²/s, 0.15 ± 0.02 µm²/s and 0.09 ± 0.01 µm²/s for early-, mid-, and late-stage respectively.

Estimation of aggresome diameter and cytoplasmic viscosity from Slimfield data

We estimated the lower limit for the aggresome diameter under early-, mid- and late-stage induction with HokB by modeling using the following method. We collated certain tracks within each early-, mid- and late-stage dataset that contained one EGFP molecule to within experimental error on the basis of the measured foci brightness values. We then generated the mean average MSD versus time interval relation across of the single EGFP tracks found after 500 frames of the start of the laser illumination. We then modeled the mean average MSD for each as that of a confining circular domain of radius *r* (*34*). In a confined diffusion in a circular domain, MSD is:

$$MSD = \delta d^2(t) \geq r^2 \left(1 - 8\sum_{m=1}^{\infty} exp\left[-r_{1m}^2 \frac{t}{\tau}\right] \frac{1}{r_{1m}^2 (r_{1m}^2 - 1)}\right) (34)$$

The *d* is the coordinates of particles. In the long-time limit t »$\tau$, the MSD converges to $r^2$ (*34*). We then equated the maximum observed mean average MSD value in each dataset to an estimate for the lower limit of *r* and the lower limit of the aggresome diameter as $2r$.

Estimates for the cytoplasmic viscosity were determined by using the Stokes-Einstein equation (*35*):

$$D = \frac{kT}{6\pi\eta r} \quad (1)$$

$D$ is the diffusion coefficient from 0.29 µm$^2$/s, 0.26 µm$^2$/s and 0.19 µm$^2$/s for early-, mid-, late-stage respectively; $k_B$ is Boltzmann's constant; $T$ is the absolute temperature; $\eta$ is the dynamic viscosity; $r$ is the radius of the spherical particle (see Table S4).

Determining the Number of Proteins per Aggresome and Number of Proteins per Cell

The number of EGFP molecules per aggresome was determined by calculating the total integrated intensity in the area of each aggresome mask. Within the aggresome mask, we first subtracted the mean background level of wild type *E coli* cells (MG1655) from each pixel, and then we defined mean value intensity among all the pixels. This mean value was divided by the characteristic single fluorophore intensity value. After that, this value was multiplied by the aggresome mask area. The probability distribution for the number of proteins per aggresome of HslU-EGFP was rendered using Gaussian kernel density estimation (KDE) (*9*) (Fig1F). To calculate the number of EGFP contained in the whole of a cell (i.e. the copy number), we used a previously developed CoPro algorithm that utilizes numerical integration and convolution of pixel intensities (*33*). In brief, we can calculate the fluorophore density from each pixel in the images. The cell pool was modeled as uniform fluorescence over rod-shaped cell of various cell length (depending on cells) and 800 nm width, which consisted in 2D of a rectangle capped by a half-circle at either end (corresponding to the 2D projection of the 3D *E. coli* cell shape of a cylinder capped by two hemispheres). The auto-fluorescence background level was determined by calculating the integrated mean intensity of parental strain (MG1655). The camera detector noise background defined by randomly selected area out of cell regions (mean size of camera background area ≈ a 25×25 square ROI) was subtracted from the mean pixel intensity value.

FRAP Analysis

The intensity traces of the half-FRAP or whole-FRAP bleached region targeted by the laser bleaching pulse were recorded by ZEN Black program as a function of time. For a photobleaching correction (i.e. due to photobleaching occurring during each sampling timepoint after the original focused laser bleach), we selected three cells in each video which were not been targeted and generated the equivalent intensity trace for them. To estimate the correction factor at each time point, we calculated the total integrated pixel intensities for each cell, and normalized this by the total integrated pixel intensity for each cell for the first time point in the series. We then calculated mean normalized intensity value for each timepoint across these three cells. The correction factor was taken as the reciprocal of this mean normalized intensity at each time point. To generate a corrected intensity for the FRAP intensity values after the focused laser bleach we then multiplied these by the correction factor for each corresponding timepoint. This process was carried out in each video and then the recovery curve was normalized by the initial intensity. For a final recovery curve, all the normalized recovery curves were averaged to determine the mean values. Fluorescence loss regions in half-FRAP experiments were selected as the half-unbleached aggresome region using ImageJ and then corrected for photobleaching using the same method as above. Fluorescence loss regions in whole-FRAP experiments were extracted by masking out the bleaching region of either one aggresome at the opposite pole of the cell, or from the whole cell area out the bleach region, as appropriate, and corrected for photobleaching as above. For half-FRAP experiments, aggresomes which showed fluorescence loss from the half-unbleached region were indicative of HslU-EGFP molecule mobility between unbleached and bleached regions.

Antibiotic Sensitivity Assay

Bacterial cultures were diluted by 1:20 into fresh LB with the following concentrations of antibiotics as appropriate for each sensitivity assay: 100 μg/ml for ampicillin, 10 μg/ml for carbencillin, 150 μg/ml for amoxicillin, 1 μg/ml for ciprofloxacin, 5 μg/ml for norfloxacin and 5 μg/ml for ofloxacin, respectively. Then the culture was returned to the 37ºC shaker for another 4 hrs. Samples were withdrawn and appropriately diluted in LB medium and spotted on an LB agar plate for overnight culture at 37 ºC. Colony counting was performed on the next day.

P1 Phage Sensitivity Assay

Bacterial cultures and P1 phages were added to LB medium (supply with 5mM $CaCl_2$ and 5mM $MgCl_2$ to activate infection, MOI=100). After 10 mins incubation on ice for adsorption, the cells were transferred to 37 ºC for 20 mins to complete infection (phage DNA injection). Then sodium citrate (final concentration 10 mM) was added to the culture to deplete $Ca^{2+}$ to terminate the infection process. Bacterial cells were washed with LB supplemented with 10mM sodium citrate following centrifugation at 3,000g for 2 mins. Samples were diluted as appropriate in PBS and spotted on LB agar plates (supplemented with 10 mM sodium citrate) for overnight culturing at 37 ºC. Colony counting was performed on the following day.

Statistics

Statistical tests were performed with the commercial software packages SPSS 18.0 as indicated in the figure legends.

### Supplementary Text

Individual-based lattice model

*Summary*

We developed an individual-based lattice model that inferred parameters from the single-molecule experimental information from Slimfield microscopy to predict the collective behaviour (LLPS, FRAP) at the continuum level *(14, 17)*. The key ingredients to this model are 1) a discretization of the elongated cell geometry on a 1×3 μm$^2$ square lattice; 2) the diffusivities of- and interactions between proteins, and 3) the distinction between two classes of proteins, namely those that *drive* the formation of aggresomes by LLPS, and those that only weakly bind to the aggresomes and can serve as *probes* for aggresome formation (we interpreted the candidate aggresome biomarker proteins of EGFP labeled HslU, Kbl and AcnB as LLPS probes). In our modeling, we represented LLPS-driving proteins by a single mean-field-type protein 'A', and the probes by a protein `B'. As we show below, this distinction enabled us to describe LLPS by the number of A-proteins, $N_A$, its interaction energy, $\varepsilon_{AA}$, its diffusivity, $D_A$, but also the numerical choice of a lattice spacing, $\Delta x$, which affects the surface tension of the aggresome. For a diffusion length of $\Delta x$=4 nm, two aggresomes form in the experimentally relevant time scale of 1-2 hours with a diffusivity of $D_A$ =0.2 μm$^2$/s. Regardless of the lattice spacing, the FRAP recovery curves require small diffusivities of the order $D_B$=10$^{-4}$ -10$^{-3}$ μm$^2$/s for the probes; our simulations are carried out at a computationally more feasible $\Delta x$=20 nm and $D_A$ =6·10$^{-3}$ μm$^2$/s (details given below). As long as the number of B-proteins, $N_B$, remains sufficiently low, the FRAP curves can be described independently by a binding energy $\varepsilon_{AB}$, the diffusivities inside, $D_{B,in}$, and outside the aggresome, $D_{B,out}$, and the position and radius of the simulated laser that photobleaches the B-proteins. Below, we first detail the algorithm, then the choice of parameters to simulate LLPS, and finally the choice of parameters to simulate FRAP. Finally, we used our results to discuss the experimental variations in the FRAP curves measured for HslU, kbl and AcnB.

*Algorithm*

To model LLPS, the A-proteins were located at lattice sites and interacted through an attractive nearest-neighbour interaction energy $\varepsilon_{AA}$. In 2D, LLPS occurred when $\varepsilon_{AA}$ was larger than the critical value of approximately 1.8 $k_BT$, with $k_B$ the Boltzmann constant and $T$ the absolute temperature *(16)* (the critical value of $\varepsilon_{AA}$ depends on the type of lattice used, e.g., hexagonal, 2D, 3D, etc.: $\varepsilon_{AA}$ should be considered a qualitative tuning parameter rather than an energy that is directly related to the structure of a real protein). Just above the critical value, the domain purity is low and the interfaces are wide and phase separation is difficult to distinguish, especially in the confined cell geometry. For increasing values of $\varepsilon_{AA}$ the domain purity increased and the interfaces became sharper (the interface width is directly determined by $\Delta x$, and, like $\varepsilon_{AA}$, is dependent on the type of lattice used). Using a combined choice of the value of $\varepsilon_{AA}$, $\Delta x$ and the number of proteins A, $N_A$, the experimental features of aggresome formation can be qualitatively reproduced. That is, in a range of $\Delta x$ of 10-40 nm we found that a value $\varepsilon_{AA} = 2.2\, k_BT$ and an A-concentration of up to $100\%\times N_A / (1\times 3\ \mu m^2 / \Delta x^2)$=40% A-rich droplets in an A-poor environment were formed. In our simulations, we included $N_A$=2,350 A proteins on a 50×150 lattice (the concentration is approximately 31.3%). For <2,000 A proteins, droplets mostly grew by ripening in contrast to the experiments where droplet fusion is observed.

In addition to protein A, our simulations also included 200 B protein molecules that probe the LLPS. This number was chosen to be sufficiently small to not affect the dynamics of A, but sufficiently large to collect enough statistics. This implies that within our model differences in copy number only affect the overall stoichiometry, but does not alter the physical trends; the number of B protein molecules in the simulation need not reflect the actual experimental copy number. In order to facilitate the diffusion of probe B into A-rich droplets, protein B may occupy the same site as protein A. In fact, diffusion of B into the aggresomes is promoted by an attractive interaction energy $\varepsilon_{AB}$. The binding energy primarily affects the rate by which B may escape the aggresome and the partition of B inside, $c_{in}$, and outside, $c_{out}$, the aggresome. At thermal equilibrium the chemical potentials inside, $\mu_{in}$, and outside, $\mu_{out}$, the aggresomes are equal and given by $\mu_{in} = \Delta H + k_BT \ln c_{in} = \mu_{out} = \ln c_{out}$, with $\Delta H$ the binding enthalpy of protein B to the aggresome, so

$$\frac{c_{in}}{c_{out}} = \exp(\Delta H / k_B T) \quad (2)$$

Collecting all interaction terms, the total internal energy is given by

$$U = N_{AA}\varepsilon_{AA} + N_{AB}\varepsilon_{AB} \quad (3)$$

with $N_{AA}$ the number of nearest A-A neighbours and $N_{AB}$ the number of lattice sites shared by A and B. This interaction energy biased the diffusion dynamics, which was modeled by enabling proteins to hop to the nearest-neighbour lattice sites. Using a simple kinetic Monte Carlo algorithm ((*36*) (*14*), at every time step, with a time increment discussed below, a random protein and hop direction was selected out of a list of $N_{proc}$ potential processes (pseudo-random numbers are generated using the SIMD-oriented Fast Mersenne Twister (*37*)). If the hop was not forbidden (a site may not contain more than one A or B protein), the rate, $r$, of the process was calculated (see below). If the rate equals the maximum rate $r_{max}$ (note that $r_{max}$ corresponds to the bare hopping frequency, while interactions may only slow down the dynamics: $r \leq r_{max}$), the rate is accepted. If it is smaller, the rate is accepted but with a probability $r/r_{max}$. Regardless if a process is accepted or rejected, time is updated as

$$\Delta t = \frac{-\ln u}{N_{proc} r_{max}} \quad (4)$$

with $u$ a uniform random number, $0<u\leq 1$.

The rate of each process depends on which protein hops, as well as on the local environment of the protein. In general, the rate is given by

$$r = \nu \cdot \min\big(1, exp(-\Delta U / k_B T)\big) \quad (5)$$

according to the Metropolis algorithm. The prefactor $\nu$ is, in absence of interactions, related to a diffusivity $D$ as $\nu = D/\Delta x^2$. In our simulations with $\Delta x$=20 nm, we used $D_A = 6\cdot 10^{-3} \mu m^2/s$ to match the experimental time scale of the order of an hour to form two aggresomes (for $\Delta x$=4 nm the more realistic value of $D_A$=0.2 μm$^2$/s may be used; this would increase the simulation time by a factor >3000). The FRAP recovery curves are virtually unaffected by the lattice spacing with fixed diffusivities of B $D_{B,in}$=2 $10^{-4}$ μm$^2$/s for when B hops between lattice sites that are co-occupied by A, $D_{B,out}$=$10^{-3}$ μm$^2$/s when B hops between lattice sites that both do not contain A, and $D$=($D_{B,in}$ + $D_{B,out}$ )/2 if one of the sites contain A.

*Modelling of LLPS*

We quantified the progress of LLPS using a characteristic length scale, $R$ ($t$), obtained from the structure factor *(14)*. The structure factor, $S$ ($q$), with $q$ the wavenumber, was obtained by taking the Fourier transform of the configuration of the A proteins (visualised by the images in Fig. 3 and Fig. S4) and taking the angularly averaged square of the Fourier transform. The structure often had a low signal-to-noise ratio, and a numerically more robust value was obtained using the inverse Fourier transform, which is the correlation function, $C(r)$ as a function of the distance $r$. In line with our previous work *(14)* we used the first minimum of $C(r)$ to define the characteristic length scale. The measures $S(q)$ and $C(r)$ were displayed in Fig. S4A and S4B, respectively, for the simulation shown in Fig 3A in the main text.

To collect statistics, we ran every simulation for various random number seeds. While these may lead to qualitatively different late-stage (at the stage the droplet size approaches the size of the cell) structures, see Fig. S4C, for early times we expected the length scale to increase according to the usual relation $R(t)$–$R_0$ ~ $t^{1/3}$ characteristic for Ostwald ripening and Brownian coalescence (*18*). Indeed, in Fig S4D we averaged this quantity (with $R_0$=0.03, 0.06 and 0.12 μm for Δ$x$=10, 20, 40 nm, respectively) over 10 simulations and plotted it against time. We find that $R(t) - R_0 \propto (D_A\, t/\Delta\, x^2)^{1/3}$ up to a plateau where typically two aggresomes are formed. We have verified that the addition or removal of the B-proteins (up to 2.1 v%; well above the experimental concentration) did not significantly affect this growth curve; neither did the presence of an excluded volume region that may represent the nucleoid (Movie S3). The plateau is reached after approximately an hour when $D_A$=6·$10^{-3}$ μm$^2$/s and Δ$x$=20 nm. We extrapolate that the experimental value $D_A$=0.2 μm$^2$/s may be used when the lattice spacing is as small as Δ$x$=4 nm. This value may slightly change when different values for $\varepsilon_{AA}$ would be chosen. Furthermore, both the lattice spacing and $\varepsilon_{AA}$ alter the width of the interface and may affect the respective contributions of Ostwald ripening and droplet fusion. Fortunately, the FRAP curves are unaffected by the lattice spacing. Therefore, we have chosen a computationally convenient lattice spacing of Δ$x$=20 nm with a low A diffusivity of $D_A$ =6·$10^{-3}$ μm$^2$/s. For these values, the qualitative features of the aggresomes (Fig. S4C) and the time scale of aggresome formation (Fig. S4D) are in agreement with the experiments.

*Modeling of FRAP*

Motivated by our assumption that the dynamics of LLPS and fluorescent recovery could be decoupled, we set up idealized simulations where protein B can diffuse and interact with A-proteins whose spatial coordinates are kept constant to fix the aggresomes centres at 500 nm from the respective poles and their radius fixed at 400 nm (see Fig. S5A). While this size is somewhat larger than in the experiments and simulations with mobile A proteins, it does not affect the early stages of half-FRAP. We do expect that smaller aggresome sizes lead to a larger amplitude of whole-FRAP recovery. In Fig. S5B we showed reasonable consistency of the simulated FRAP curves between a simulation with Δ$x$=10 nm and Δ$x$=20 nm. In these simulations, we set the outside diffusivity to $D_{B,out}$=$10^{-3}$ μm$^2$/s, as this gave the correct order of magnitude in the recovery time, and manually tuned the inside diffusivity, $D_{B,in}$, binding energy, $\varepsilon_{AB}$, and laser radius, $R$ to match the experimental data. The fact that these values are much smaller than the mean values for the diffusivity measured (Fig 2B) suggests that the low-mobility tail of the distribution, possibly due to heterogeneity in the cytoplasm, determines the recovery on a macroscopic scale. We emphasize that a change in the choice of $D_{B,out}$ as well as the position of the laser in the half-FRAP simulations affects the final values. In these simulations we used a lattice spacing $\Delta x$=10 nm.

In Fig S5B we investigated the influence of the laser radius on the FRAP transients. While the half-FRAP curve was barely affected by variations of 380-400 nm, we found there was a narrow window just below 400 nm where the best match between whole-FRAP simulation and experiment were obtained. This radius, which was slightly smaller than the radius of the simulated aggresome, left some proteins at the rim of the aggresome unbleached and enabled them to govern fast, but low intensity, recovery. The need of a slightly smaller focus radius than the aggresome radius seems general in our simulations, and was also observed when LLPS and FRAP are simultaneously simulated. We speculate in the experiments the broad range of diffusivities also enables a small fraction of proteins to swiftly diffuse towards the aggresome and lead to fast initial recovery, and the sensitivity of FRAP signal on the laser focus may be weaker.

In Fig S5C we varied the binding energy, $\varepsilon_{AB}$, in the range 1.8-2.6 $k_BT$. In this range, the amplitude of the whole-FRAP curve monotonically decreased with an increasing binding energy due to an increasing concentration of B proteins inside the aggresome (see Eq. 2). At a value of approximately 2.5 $k_BT$ the best agreement with the experimental whole-FRAP curve was found. - For the simulations where LLPS and FRAP were simultaneously modeled (i.e., with mobile A-proteins, see Fig. 3 in the main text) a value 2.2 $k_BT$ gave a better match between the simulation and experiment. This suggests that the radius of the aggresomes and impurities (protein A present outside the aggresome, and vacant lattice sites inside the aggresome) affect the precise value of the binding energy. Indeed using the experimental inside and outside concentrations (see Table S4), Eqn. 2 suggests a binding enthalpy of approximately 1.4 $k_BT$.

In Fig. S5D we varied the diffusivity of protein B inside the aggresome in the range 1-5×$10^{-4}$ μm$^2$/s while keeping the outside diffusivity constant at $10^{-3}$ μm$^2$/s. In this case, the whole FRAP curve remains virtually unaffected, while the half FRAP recovery shift on the time axis proportional to the value of $D_{B,in}$. Using these simulations, we estimated the inside diffusivity to be approximately 2×$10^{-4}$ μm$^2$/s. Like the value of the outside diffusivity, this rather small value suggests that heterogeneity/distribution in mobility in the cytoplasm limits the macroscopic recovery dynamics.

In summary, for a fixed bleaching position and radius, the simulated FRAP curves can be parameterized using the diffusivity of the probe protein inside and outside the aggresome, and the binding energy. In terms of these parameters we then interpreted the variations in the FRAP curves measured for the all three proteins HslU, Kbl and AcnB shown in Fig. S5E. Given that the half-FRAP curves are predominantly determined by the inside diffusivity, and consequently have a ~$t^{1/2}$ time dependence, we shift the time axis such that the half-FRAP data collapses onto a master curve FRAP=$At^{1/2}$. We remark that the whole-FRAP recovery is achieved by a combination of diffusion inside the aggresome, exchange of proteins with the cytoplasm at short ranges and exchange of proteins between the two aggresomes at a long range: the apparent $t^{0.21}$ power law fitted to the whole-FRAP data is expected to be a purely heuristic effective fit to the model for all these processes combined. Figs. S5F-S5H show that after collapsing the half-FRAP data onto the master curve, the whole-FRAP data is now only affected by the outside diffusivity and by the binding energy. Indeed, if the outside diffusivity is low, then protein transport to the bleached area is small and the vertical offset of the whole FRAP curve decreases. If the binding energy is high, the protein concentration outside the aggresomes is low and the whole-FRAP signal also has a lower amplitude.

By applying this method to the experimental data we have obtained Fig S6H. Collapsing the half-FRAP data onto the master curve yielded estimates for the inside diffusivities of Kbl and AcnB to be 0.8 and 0.4 times the inside diffusivity of HslU, respectively. Further, the whole FRAP intensity decreased from Kbl to HslU to AcnB. The differences in the protein diffusivities may originate from differences in size and (energetic or hydrophobic) interactions with its environment. Within our model, we interpret that this decreased recovery rate originates from a lower concentration and/or a lower diffusivity of the respective proteins outside the aggresomes.

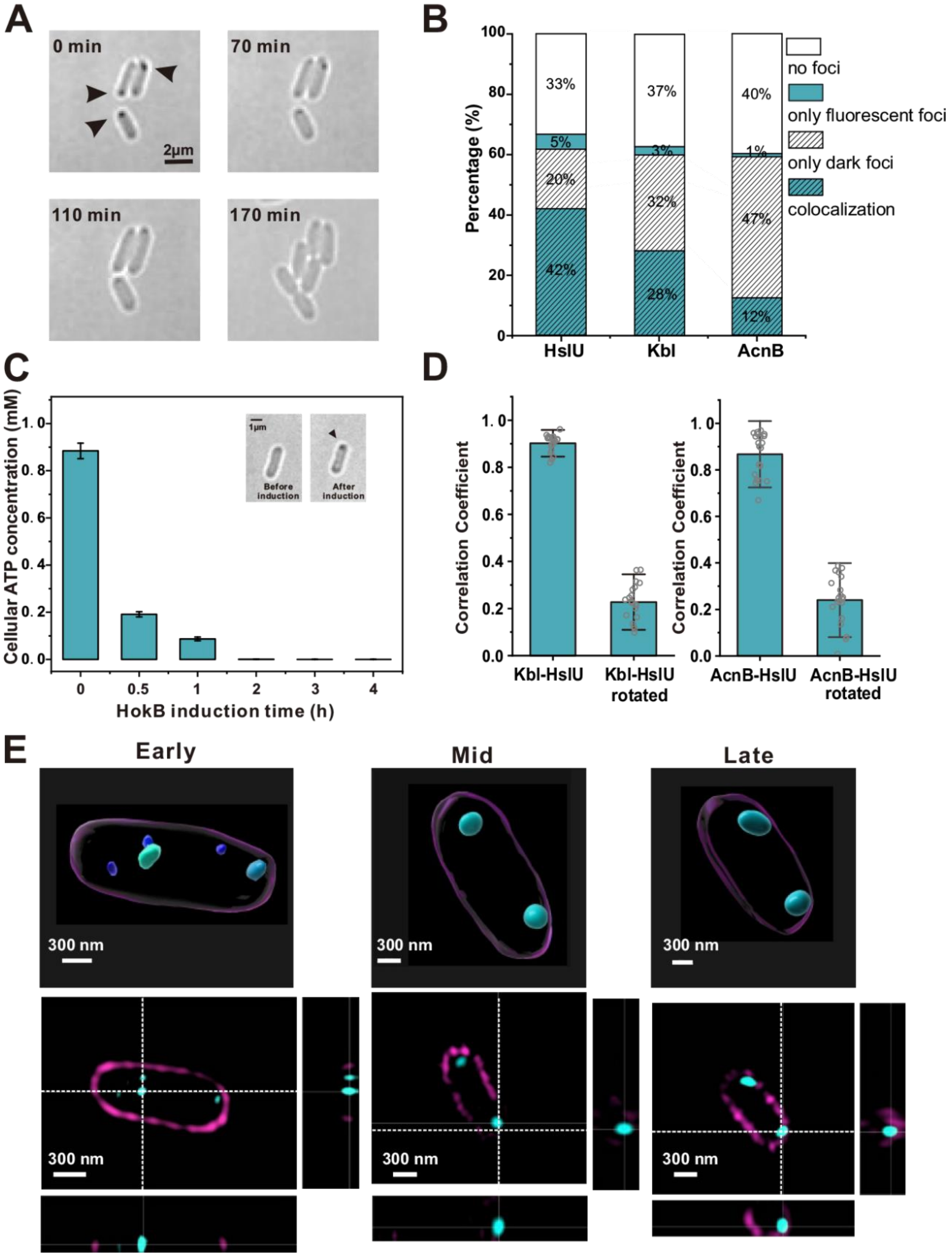


**Fig. S1. Inducing HokB to stimulate aggresome formation.**

(A) Brightfield images showing disassemble of ATP-dependent aggresome (black arrow) when cells experience fresh LB media. (B) Colocalization degree of brightfield dark foci and fluorescent foci of HslU-EGFP, Kbl-EGFP and AcnB-EGFP after 24 hrs culture. (C) Cellular ATP concentration as a HokB induction time. (Insert: brightfield images of the cell before and after HokB induction, error bar indicates SD.) (D) Colocalization analysis of HslU-mCherry & Kbl-EGFP fluorescent foci and HslU-mCherry & AcnB-EGFP fluorescent foci after 5 hrs HokB induction (only cells with fluorescent foci in both mCherry and EGFP channels were analyzed, N=20 for each group, error bar indicates SD). (left) Pearson's correlation coefficients of HslU-mCherry fluorescent foci and Kbl-EGFP fluorescent foci of cells overexpressing HokB before and after rotating HslU-mCherry image by 90 degrees. (right) Pearson's correlation coefficients of HslU-mCherry fluorescent foci and AcnB-EGFP fluorescent foci of cells overexpressing HokB before and after rotating HslU-mCherry image by 90 degrees. (E) SIM images of typical cells during aggresome formation (cyan: HslU-EGFP, magenta: cell membrane stained by FM4-64). Top: 3D view rendering images. Bottom: orthogonal view images.

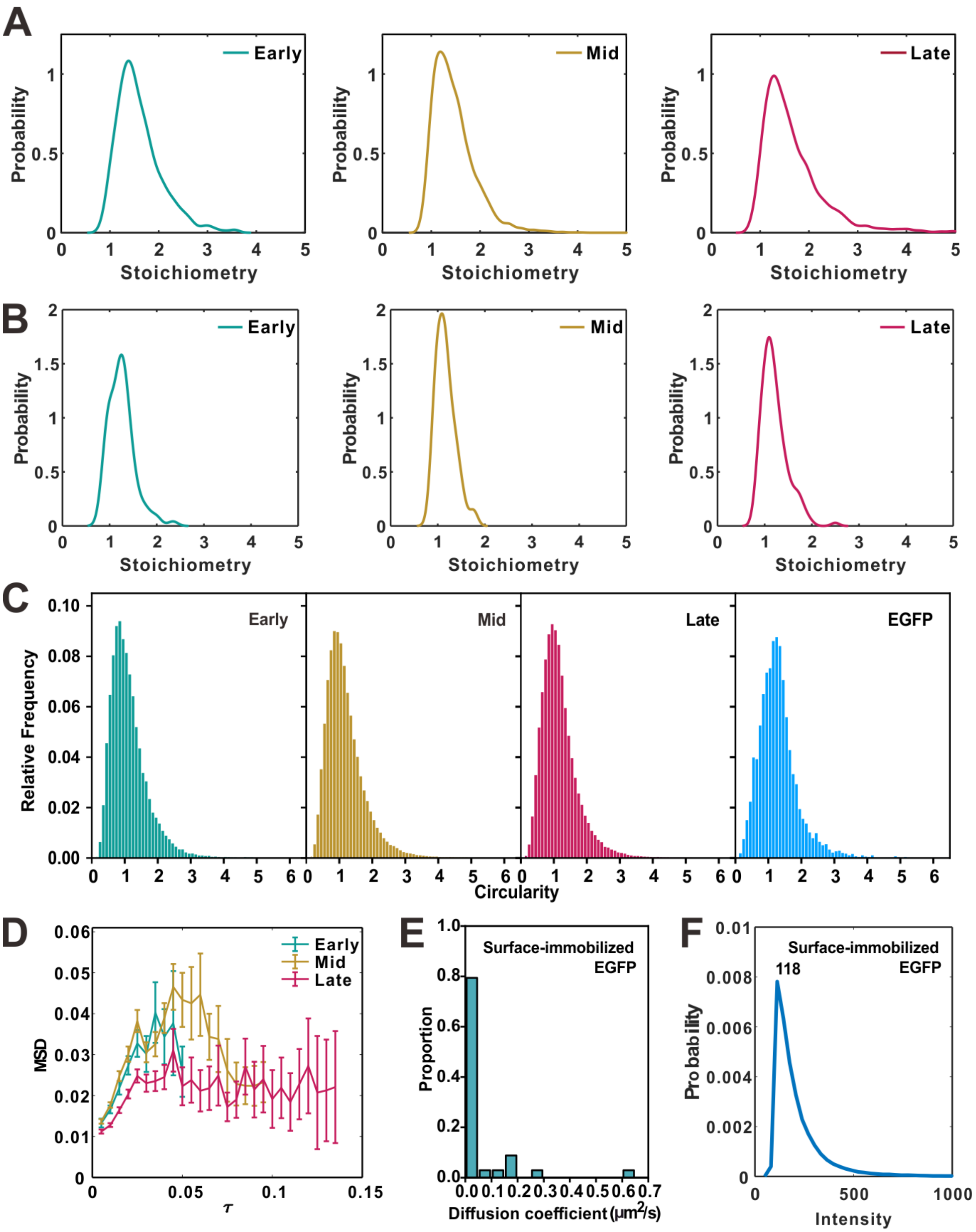


**Fig. S2. HslU mobility and circularity**

(A) Kernel density plots of stoichiometry of HslU-EGFP per focus from tracks found from 500 frames until the end of imaging at different HokB induction stages. (B) Kernel density plots of stoichiometry of HslU-EGFP per focus from tracks found at the end of photobleach process (from 1500 frames of the start of laser illumination until the end) at different HokB induction stages. All the peaks are around 1, which confirmed that the tracks for defining *Dg* are single molecules. (C)

Circularity of single-molecule spots and surface-immobilized EGFP. (D) Time versus MSD relations at the different HokB induction stages. The maximum observed mean average MSD values in the plots were used for defining diameters of aggresomes at the different HokB induction stages. (E) Histogram of the diffusion coefficients of surface-immobilized EGFP (number of molecules N=34). (F) The value of single intensity (118) is the characteristic intensity of surfaced-immobilized EGFP molecule.

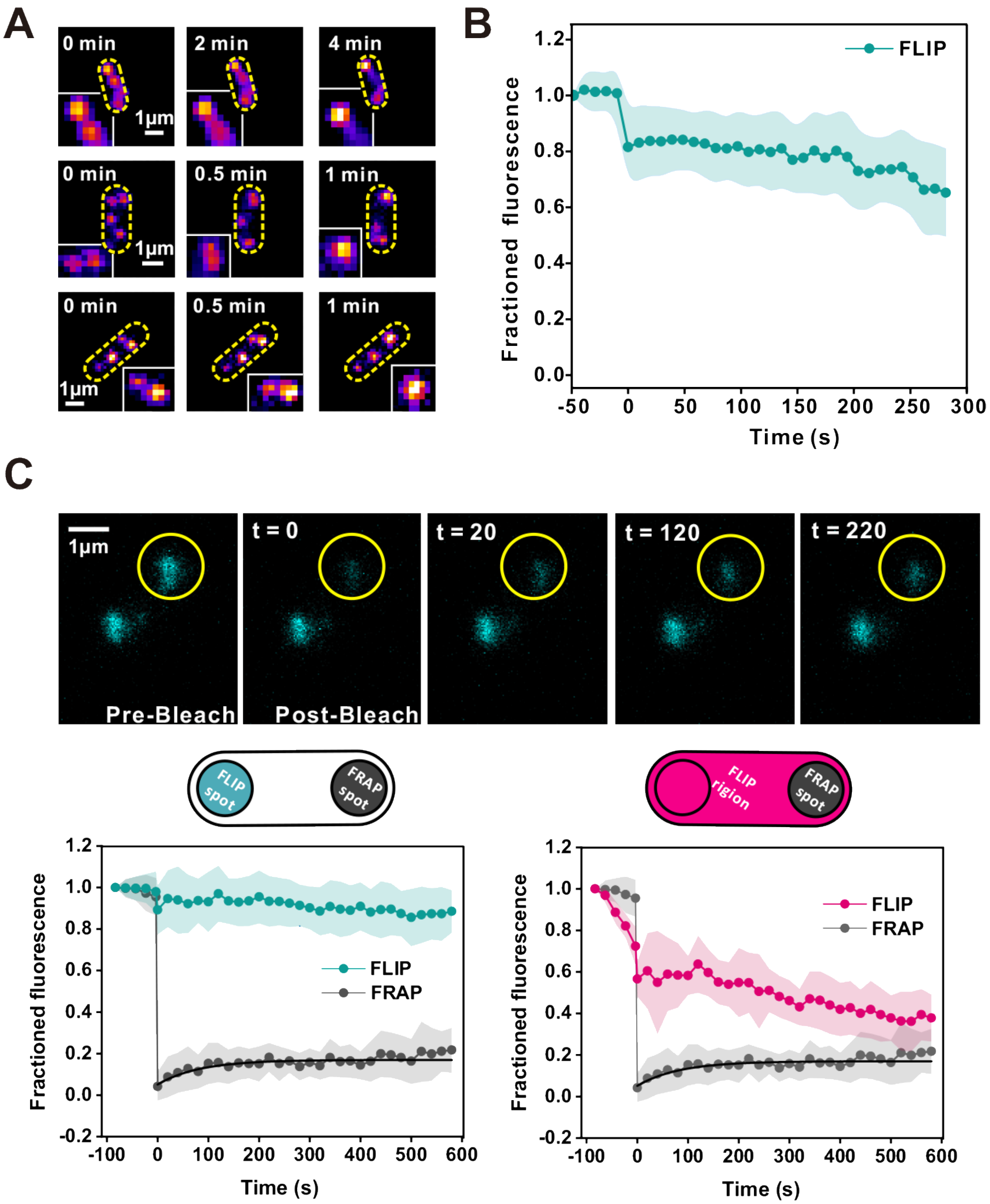


**Fig. S3. HslU turnover.**

(A) Fluorescence images of two aggresomes fusing. (B) Circularity of the aggresome at the different stages. Early: 0.96 ± 0.06, Mid: 1.07 ± 0.01, Late: 1.03 ± 0.01 (C) Mean half-FLIP trace (number of aggresomes N=29). The focused laser has a diameter of approximately 0.65 μm, so on average the cytoplasmic pool of bleached HslU-EGFP for a typical 3 μm long cell is ~1/4 whereas the cytoplasmic pool of unbleached HslU-EGFP is ~3/4 of the total cell area. If there is turnover between cytoplasm and aggresome (as we observe from whole-FRAP experiments) then a larger unbleached EGFP pool increases the apparent net rate of recovery of intensity in half-FRAP compared to the apparent rate of loss of intensity in half-FLIP. (D) An example whole-FRAP experiment of an aggresome; (top five images) fluorescence images showing fluorescence recovery from the bleached aggresome over time, half-time recovery constant 32 ± 18 secs. (Left bottom) Mean fluorescence recovery curve from the bleached aggresome and mean fluorescence loss from a second aggresome in the cell for cells that only contained two detectable aggresomes. (Right bottom) Mean fluorescence recovery curve from the bleached aggresome and mean fluorescence loss from all the entire cell area apart for the bleached aggresome.

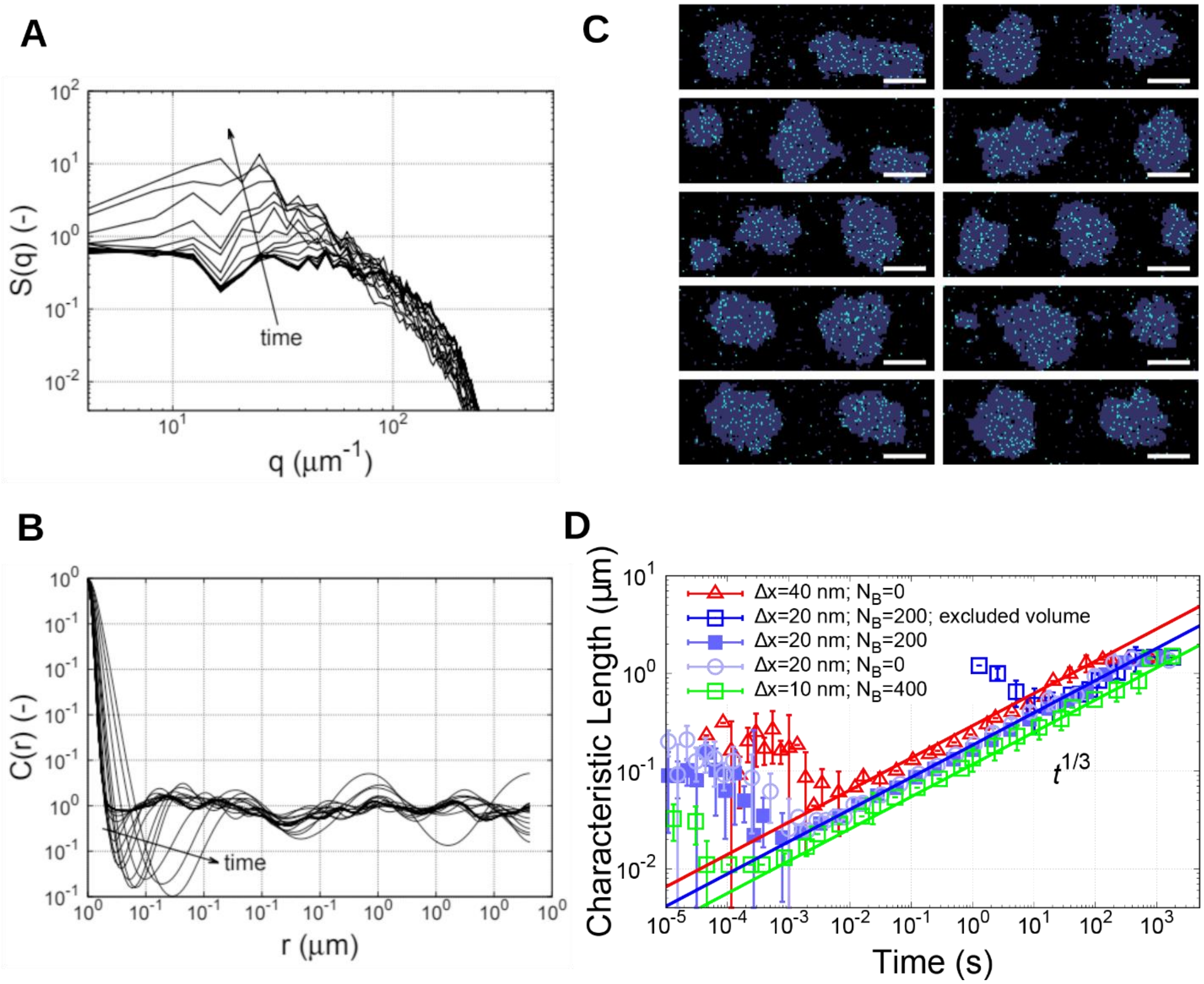


**Fig. S4. Quantification of LLPS**

(A) The structure factor, *S(q)*, against the wavenumber, q, as calculated from the configuration of protein A in the morphology images in Fig. 3 in the main text. As time proceeds, a peak emerges that shifts to decreasing wavenumbers. (B) The correlation function, *C(r)* against length scale, r, as determined from the structure factor in panel A. The first minimum is used as the characteristic length scale of the morphology. (C) Variations in morphology after $10^9$ time steps (approximately 8,500 seconds of simulated real time) for 10 different random number seeds. Typically, after this time two or three droplets have formed. The scale bar represents 500 nm. (D) The characteristic length scale is plotted against time for simulations with and without proteins B present and for two lattice spacings. The influence of protein B on LLPS is negligible. The lattice spacing does affect the rate of LLPS due to its influence on the surface tension.

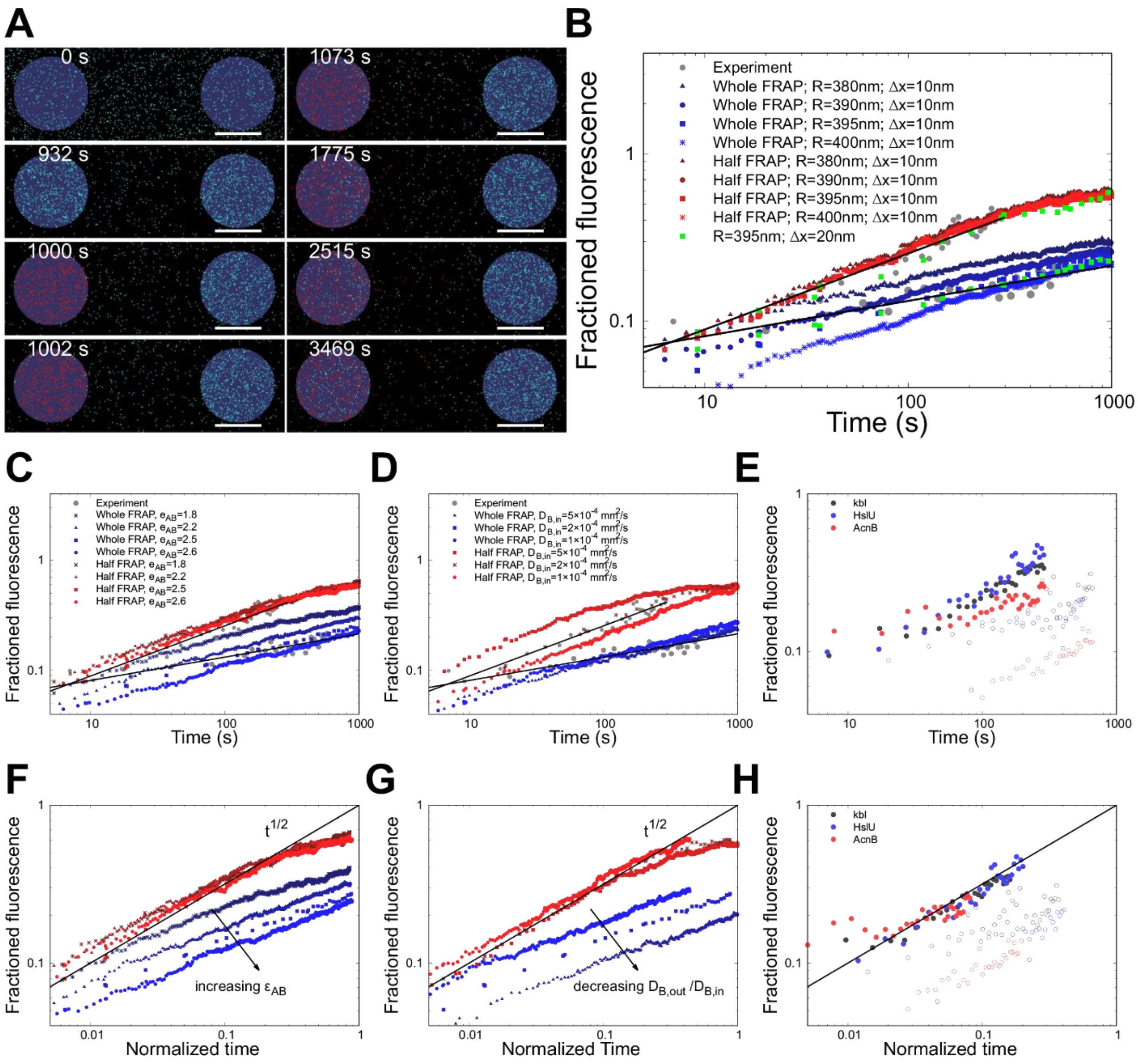


**Fig. S5 Exploring parameter space in simulations.**

The influence of the inside, $D_{\mathrm{B,in}}$, and outside diffusivity, $D_{\mathrm{B,out}}$, and the binding energy, $\varepsilon_{\mathrm{AB}}$, on FRAP transients is investigated. (A) The diffusion of protein B is simulated while protein A is fixed in perfectly round aggresomes with a 400 nm radius. At $t$=0 B is randomly distributed; after being equilibrated for 1000 seconds protein B is bleached within a simulated focus. The scale bar represents 500 nm. (B) Variations of the radius of the laser focus predominantly affected whole FRAP transients, while the half-FRAP transients were insensitive to this variation. (C-D) For a fixed laser position, the binding energy (C) and inside diffusivity (D) are varied, which predominantly affected the whole and the half FRAP data, respectively. (E) The experimental differences in FRAP transients between the three proteins klb, HslU and AcnB; the closed and open circles represent half- and whole-FRAP measurement, respectively. (F-H) To interpret the experimental data of (E), the time scale of (C-D) is normalized by collapsing the half FRAP data onto the master curve FRAP=$A \cdot t^{1/2}$. The prefactor $A$ provides values for the diffusivities of the proteins inside the aggresome. The remaining variations in the whole FRAP data are interpreted to originate from a combination of differences in the outside diffusivity and binding energy.

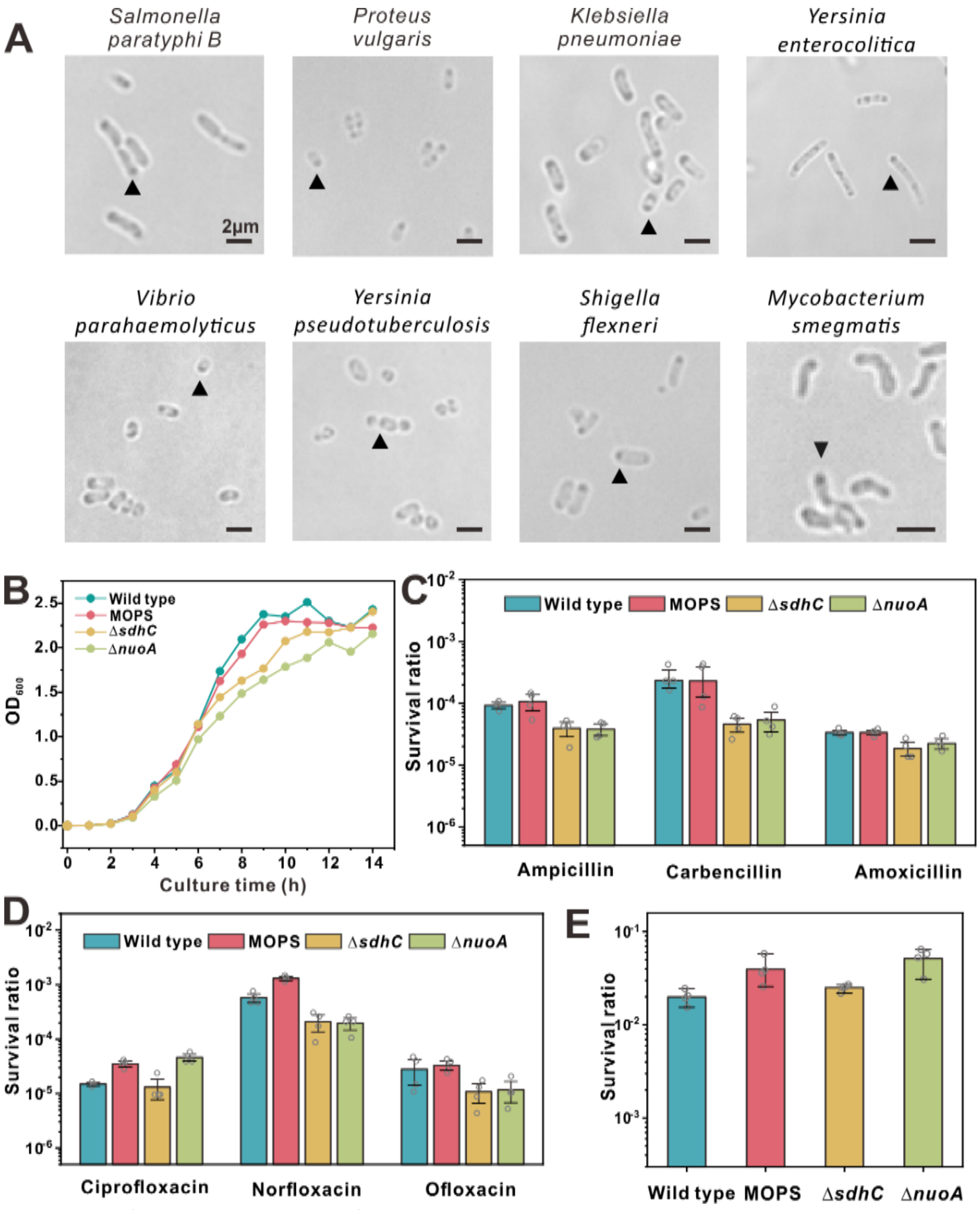


**Fig S6. Role of LLPS in bacterial fitness.**

(A) Brightfield images of different Gram-negative bacterial species in late stationary phase. Aggresome formation (black arrow) is widely observed. (B) Growth curve of different strains (wild type, MOPS, Δ*sdhC* and Δ*nuoA*). (C) Cell survival rate after 4 hrs β-lactam antibiotic treatment of different strains cultured in LB for 12 hrs. (D) Cell survival rate after 4 hrs fluoroquinolone antibiotic treatment of different strains cultured in LB for 12 hrs. (E) Cell survival rate after P1 phage infection (MOI=100) of different strains cultured in LB for 12 hrs (error bar indicates SE).

**Table S1. Strains used in this study.**

| Bacterial Strains | Source | Genotype | Doubling time (min) |
|---|---|---|---|
| MG1655 | Yale Genetic Stock Center(CGSC#:6300) | Wild-type | 23.5±0.7 |
| HslU-EGFP | LAB strain | MG1655 *ΔaraD-B* hslU::EGFP | 23.4±0.8 |
| Kbl-EGFP | LAB strain | MG1655 *ΔaraD-B* kbl::EGFP | 24.0±1.6 |
| AcnB-EGFP | LAB strain | MG1655 *ΔaraD-B* acnB::EGFP | 23.6±0.8 |
| Kbl-EGFP & HslU-mCherry | LAB strain | MG1655 *ΔaraD-B* kbl::EGFP hslU::mCherry | 25.9±1.3 |
| AcnB-EGFP & HslU-mCherry | LAB strain | MG1655 *ΔaraD-B* acnB::EGFP hslU::mCherry | 25.7±1.2 |
| *ΔnuoA* | LAB strain | MG1655 *ΔnuoA* | 22.7±1.5 |
| *ΔsdhC* | LAB strain | MG1655 *ΔsdhC* | 23.4±0.5 |
| *ΔnuoA* & HslU-EGFP | LAB strain | MG1655 *ΔaraD-B ΔnuoA* hslU::EGFP | 23.5±1.2 |
| *ΔsdhC* & HslU-EGFP | LAB strain | MG1655 *ΔaraD-B ΔsdhC* hslU::EGFP | 24.1±0.8 |

**Table S2. Plasmids used in this study.**

| Name | Gene | Resistance |
|---|---|---|
| pBAD-hokB | araBAD-hokB | Chloramphenicol |
| pSIM6 | (*28*) | Ampicillin |
| pCP20 | (*38*) | Ampicillin |

**Table S3. Primers used in this study.**

| Name | Sequence (5'-3') |
|---|---|
| araB-D knockout F | GTTTCTCCATACCCGTTTTTTTGGATGGAGTGAAACGATGATTCCGGGGATCCGTCGACC |
| araB-D knockout R | GCTGTGGTTTTATACAGTCATTACTGCCCGTAATATGCCTTTGTAGGCTGGAGCTGCTTCG |
| hslU+EGFP homology F | TGCGTTGGTGGCAGATGAAGATCTGAGCCGTTTTATCCTAGGTGGATCCGGCGGTTCTGTGAG |
| hslU+EGFP homology R | TTCAGCCCCATCAAACAATGATGAAAATGATTGAACGCGAGCTACCGCCACTGCCACCGCTC |
| hslU+EGFP P1 | GGCGACGGCCCCGTGCTGCT |
| hslU+EGFP P2 | GAACGCGATTACTTGTACAGCTCGTCCA |
| hslU+EGFP P3 | ACAAGTAATCGCGTTCAATCATTTTCAT |
| hslU+EGFP P4 | CAGCAAAGGCGAGGGGGAGG |
| acnB+EGFP homology F | CACCGAGAAAGCCGATGGGGTGATTTTCCAGACTGCGGTTGGTGGATCCGGCGGTTCTGTGAG |
| acnB +EGFP homology R | GGGCATTGTGTCGTTTATGCGCAGCGCGTGCGCTGACTTTGCTACCGCCACTGCCACCGCTC |
| acnB+EGFP P1 | GGCGACGGCCCCGTGCTGCT |
| acnB+EGFP P2 | CTGACTTTTTACTTGTACAGCTCGTCCA |
| acnB+EGFP P3 | ACAAGTAAAAAGTCAGCGCACGCGCTGC |
| acnB+EGFP P4 | CCGGCAATGCACCGAAAT |
| kbl+EGFP homology F | AGCATTTACGCGTATTGGTAAACAACTGGGCGTTATCGCCGGTGGATCCGGCGGTTCTGTGAG |
| kbl +EGFP homology R | CTTCCGCTTTCAGTTTGGATAACGCTTTCATCTCACATCCGCTACCGCCACTGCCACCGCTC |
| kbl+EGFP P1 | GGCGACGGCCCCGTGCTGCT |
| kbl+EGFP P2 | TCACATCCTTACTTGTACAGCTCGTCCA |
| kbl+EGFP P3 | ACAAGTAAGGATGTGAGATGAAAGCGTT |
| kbl+EGFP P4 | CGACCACTCATCCCAGTT |

| | |
|---|---|
| Cat(HslU)-F | CACCATCGAAGAATTAAGCTACAAAGCGTAAGGATCTCCCTCTAGAGCGACGCCAGACGG |
| Cat(HslU)-R | TTCAGCCCCATCAAACAATGATGAAAATGATTGAACGCGATTACGCCCCGCCCTGCCACT |
| HslU-F | ATGTCTGAAATGACCCCACGC |
| HslU-R | GAGCCACCTAGGATAAAACGGCTCAGATCTTCATCTGC |
| Linker-mCherry-F | TTATCCTAGGTGGCTCTGGTGGCGGTTC |
| mCherry-R | TTACTTGTACAGCTCGTCCATGCCG |
| Homo-HslU-F | CACCATCGAAGAATTAAGCTACAAAGCGTAAGGATCTCCCATGTCTGAAATGACCCCACG |
| Homo-mCherry-R | TTCAGCCCCATCAAACAATGATGAAAATGATTGAACGCGATTACTTGTACAGCTCGTCCATGCCG |
| sdhC knockout F | CCCAGGGAATAATAAGAACAGCATGTGGGCGTTATTCATGTCTAGAGCGACGCCAGACGG |
| sdhC knockout R | CCTAATGCGGAGGCGTTGCTTACCATACGAGGACTCCTGCTGTAGGCTGGAGCTGCTTCG |
| nuoA knockout F | GAGCAGTGAATCTGGCGCTACTTTTGATGAGTAAGCAATGATTCCGGGGATCCGTCGACC |
| nuoA knockout R | CCATCTTAATGCCTCGCGGTTAGCGTTGACGATTAGCGATTGTAGGCTGGAGCTGCTTCG |

**Table S4. Information of aggresomes.**

Number of HslU-EGFP molecules per cell at different incubation stages, with estimates of aggresome volume based on measurements of aggresome diameter from MSD analysis and assuming a spherical shape. Errors indicated are standard deviation, number of cells measured in range 31-209. Aggregation enthalpy ($\Delta H$) of HslU in the early, mid and late stages. The aggregation enthalpies are calculated using Eq. (2), where the outside concentration is estimated as $c_{\text{out}}=(N_{cell}-N_{agg})/(V_{cell}-V_{agg})$.

| | Early | Mid | Late |
|---|---|---|---|
| Number of aggresome per cell | 2.0 | 2.4 | 2.3 |
| Number of protein per cell | 276 ± 253 | 510 ± 345 | 681 ± 437 |
| Number of protein per aggresome | 35 ± 20 | 50 ± 32 | 63 ± 35 |
| Mean cytoplasmic viscosity (cP) | 6.2 | 8.3 | 16.6 |
| $c_{in}/c_{out}$ | 4.27 ± 0.42 | 5.11 ± 1.26 | 3.78 ± 0.11 |
| $\Delta H/k_BT$ | 1.45 ± 0.10 | 1.53 ± 0.25 | 1.33 ± 0.03 |
| Volume of aggresome (μm$^3$) | 0.050 ± 0.008 | 0.055 ± 0.008 | 0.046 ± 0.009 |

**Movie S1**

**Half FRAP imaging.** By focusing a laser laterally offset approximately 0.5 µm from the center of an aggresome it was possible to photobleach approximately one half while leaving the other half intact, which we denote as "half-FRAP". We then measured the aggresome fluorescence intensity at 10 secs intervals for up to several hundred seconds after the focused laser bleach. (White dot: Bleached half-aggresome.)

**Movie S2**

**Half FRAP simulation.** The size of the cell is 1×3 µm and the volume fraction of LLPS-driving protein A is 31% (grey) and LLPS-probing protein B (unbleached: cyan; bleached: red) is 2.7% In the simulations, the aggresomes emerge and coarsen for 158 minutes, after which protein B is bleached (colored red) within a simulated laser focus region.

**Movie S3**

**Simulation with excluded volume.** Aggresome formation simulation with excluded volume in the middle of the cell. The size of the cell is 1×3 µm and the volume fraction of LLPS-driving protein A is 31% (grey) and LLPS-probing protein B (cyan) is 2.7%. The time scales of coarsening are similar to simulations without an excluded volume region.

**Supplementary references：**


27. K. A. Datsenko, B. L. Wanner, One-step inactivation of chromosomal genes in *Escherichia coli* K-12 using PCR products. *Proc. Natl. Acad. Sci. U. S. A.* **97**, 6640-6645 (2000).
28. S. Datta, N. Costantino, D. L. Court, A set of recombineering plasmids for gram-negative bacteria. *Gene* **379**, 109-115 (2006).
29. M. J. Giacalone *et al.*, Toxic protein expression in *Escherichia coli* using a rhamnose-based tightly regulated and tunable promoter system. *Biotechniques* **40**, 355-364 (2006).
30. T. Baba et al., Construction of Escherichia coli K-12 in-frame, single-gene knockout mutants: the Keio collection. *Mol. Syst. Biol.* **2**, 0008 (2006).
31. S. Bolte, F. P. Cordelieres, A guided tour into subcellular colocalization analysis in light microscopy. *J. Microsc. (Oxford, U. K.)* **224**, 213-232 (2006).
32. A. J. M. Wollman, M. C. Leake, Millisecond single-molecule localization microscopy combined with convolution analysis and automated image segmentation to determine protein concentrations in complexly structured, functional cells, one cell at a time. *Faraday Discuss.***184**, 401-424 (2015).
33. H. Miller, Z. Zhou, A. J. M. Wollman, M. C. Leake, Superresolution imaging of single DNA molecules using stochastic photoblinking of minor groove and intercalating dyes. *Methods* **88**, 81-88 (2015).
34. T. Bickel, A note on confined diffusion. *Phys. A.* **377**, 24-32 (2007).
35. T. Kalwarczyk, M. Tabaka, R. Holyst, Biologistics-Diffusion coefficients for complete proteome of Escherichia coli. *Bioinformatics* **28**, 2971-2978 (2012).
36. A. P. J. Jansen, An introduction to kinetic Monte Carlo simulations of surface reactions, (Springer-Verlag, Berlin Heidelberg, 1st edn, 2012).

37. M. Saito, M. Matsumoto, “SIMD-oriented Fast Mersenne Twister: A 128-bit pseudorandom number generator.” in *Monte Carlo and Quasi-Monte Carlo Methods* (Springer, 2008), pp. 607-622.
38. PP Cherepanov, W. Wackernagel, Gene disruption in *Escherichia coli*: TcR and KmR cassettes with the option of Flp-catalyzed excision of the antibiotic-resistance determinant. *Gene* **158**, 9–14(1995).
